# The protective effect of sickle cell haemoglobin against severe malaria depends on parasite genotype

**DOI:** 10.1101/2021.03.30.437659

**Authors:** Gavin Band, Ellen M. Leffler, Muminatou Jallow, Fatoumatta Sisay-Joof, Carolyne M. Ndila, Alexander W. Macharia, Christina Hubbart, Anna E. Jeffreys, Kate Rowlands, Thuy Nguyen, Sonia M. Goncalves, Cristina V. Ariani, Jim Stalker, Richard D. Pearson, Roberto Amato, Eleanor Drury, Giorgio Sirugo, Umberto D’Alessandro, Kalifa A. Bojang, Kevin Marsh, Norbert Peshu, David J. Conway, Thomas N. Williams, Kirk A. Rockett, Dominic P. Kwiatkowski

**Affiliations:** Wellcome Centre for Human Genetics, University of Oxford, Oxford, UK; Wellcome Sanger Institute, Hinxton, Cambridge, UK; Department of Human Genetics, University of Utah, Salt Lake City, Utah 84112-5330; Medical Research Council Unit The Gambia at the London School of Hygiene and Tropical Medicine, Atlantic Boulevard, Fajara, The Gambia; KEMRI-Wellcome Trust Research Programme, CGMRC, PO Box 230-80108, Kenya; Centre for Global Health Tropical Medicine, University of Oxford, Oxford, UK; London School of Hygiene and Tropical Medicine, Keppel Street, London, UK; Department of Infectious Diseases, Imperial College Faculty of Medicine, London W21NY, United Kingdom; MRC Centre for Genomics and Global Health, Big Data Institute, Old Road Campus, Oxford OX3 7LF, UK

## Abstract

Host genetic factors can confer resistance against malaria, raising the question of whether this has led to evolutionary adaptation of parasite populations. In this study we investigated the correlation between host and parasite genetic variation in 4,171 Gambian and Kenya children ascertained with severe malaria due to *Plasmodium falciparum*. We identified a strong association between sickle haemoglobin (HbS) in the host and variation in three regions of the parasite genome, including nonsynonymous variants in the acyl-CoA synthetase family member *PfACS8* on chromosome 2, in a second region of chromosome 2, and in a region containing structural variation on chromosome 11. The HbS-associated parasite alleles are in strong linkage disequilibrium and have frequencies which covary with the frequency of HbS across populations, in particular being much more common in Africa than other parts of the world. The estimated protective effect of HbS against severe malaria, as determined by comparison of cases with population controls, varies greatly according to the parasite genotype at these three loci. These findings open up a new avenue of enquiry into the biological and epidemiological significance of the HbS-associated polymorphisms in the parasite genome, and the evolutionary forces that have led to their high frequency and strong linkage disequilibrium in African *P. falciparum* populations.

## Main text

Malaria can be viewed as an evolutionary arms race between the host and parasite populations. Human populations in Africa have acquired a high frequency of sickle haemoglobin (HbS) and other erythrocyte polymorphisms that provide protection against the severe symptoms of *Plasmodium falciparum* ^1,2^ infection, while *P. falciparum* populations have evolved a complex repertoire of genetic variation to evade the human immune system and to resist antimalarial drugs ^3,4^. This raises the basic question: are there genetic forms of *P. falciparum* that can overcome the human variants that confer resistance to this parasite?

To address this question, we analysed both host and parasite genome variation in samples from 5,096 Gambian and Kenyan children with severe malaria due to *P. falciparum* (**Supplementary Figure 1-2** and **Methods**). All of the samples were collected over the period 1995-2009 as part of a genome-wide association study (GWAS) of human resistance to severe malaria that has been reported elsewhere^2,5,6^. In brief, we sequenced the *P. falciparum* genome using the Illumina × Ten platform using two approaches based on sequencing whole DNA and selective whole genome amplification^7^. We used an established pipeline ^8^ to identify and call genotypes at over 2 million single nucleotide polymorphisms (SNPs) and short insertion/deletion variants across the Pf genome in these samples (**Methods**). The following analysis is based on 4,171 samples that had high quality data for both parasite and human genotypes and were not closely related, of which a subset of 3,346 had human genome-wide genotyping available. We focussed on a set of 51,225 biallelic variants in the *P.falciparum* genome that passed all quality control filters and were observed in at least 25 infections in this subset. Our analyses exclude mixed genotype calls that arise in malaria when a host is infected with multiple parasite lineages. Full details of our sequencing and data processing can be found in **Supplementary Methods**.

We used a logistic regression approach to test for pairwise association between these *P. falciparum* variants and human variants selected according to four criteria: i. known autosomal protective mutations, including HbS (within *HBB*), the common mutation that determines O blood group (within *ABO)*, regulatory variation associated with protection at *ATP2B4* ^2,5,9^ and the structural variant DUP4, which encodes the Dantu blood group phenotype ^10^; ii. variants that showed suggestive but not conclusive evidence of association with severe malaria in our previous GWAS^5^; iii. HLA alleles and additional glycophorin structural variants that we previously imputed in these samples; and iv. variants near genes that encode human blood group antigens, which we tested against the subset of *P.falciparum* variants lying near genes which encode proteins important for the merozoite stage ^11,12^, as these might conceivably interact during host cell invasion by the parasite. Although several factors could confound this analysis in principle – notably, if there were incidental association between human and parasite population structure – the distribution of test statistics suggested that our test was not affected by systematic confounding after including only an indicator of country as a covariate (**Supplementary Figure 3**), and we used this approach for our main analysis. A full list of results is summarised in **Supplementary Figure 4** and **Supplementary Table 1**.

The most striking finding to arise from this joint analysis of host and parasite variation was a strong association between the sickle haemoglobin allele HbS and three separate regions in the *P. falciparum* genome (**Supplementary Figure 4 and Figure 1**). Additional associations with marginal levels of evidence were observed at a number of other loci, including a potential association between *GCNT1* in the host and *PfMSP4* in the parasite and associations involving HLA alleles (detailed in **Supplementary Methods** and **Supplementary Table 1**), but here we focus on the association with HbS.

The statistical evidence for association at the HbS-associated loci can be described as follows, focussing on the variant with the strongest association in each region and assuming an additive model of effect of the host allele on parasite genotype **(Supplementary Table 1)**. The chr2: 631,190 T>A variant, which lies in *PfACS8*, was associated with HbS with Bayes factor (*BF*_HbS_) = 1.1 × 10^15^ (computed under a log-F(2,2) prior; **Methods**) and *P =* 4.8 × 10^−13^ (computed using a Wald test; **Supplementary Methods**). At a second region on chromosome 2, the chr2: 814,288 C>T variant, which lies in *Pf3D7_0220300*, was associated with *BF*_HbS_ = 2.4 × 10^9^ and *P =* 1.6 × 10^−10^. At the chromosome 11 locus, the chr11: 1,058,035 T>A variant, which lies in *Pf3D7_1127000*, was associated with *BF*_HbS_ = 1.5×10^17^ and *P =* 7.3×10^−12^. For brevity we shall refer to these HbS-associated loci as *Pfsa1*, *Pfsa2* and *Pfsa3* respectively, and we shall use + and – signs to refer to the alleles that are positively and negatively correlated with HbS, e.g. *Pfsa1*+ is the allele that is positively correlated with HbS at the *Pfsa1* locus. All three of the lead variants are nonsynonymous mutations of their respective genes, as are additional associated variants in these regions (**Figure 1** and **Supplementary Table 1)**.

**Figure 1:**
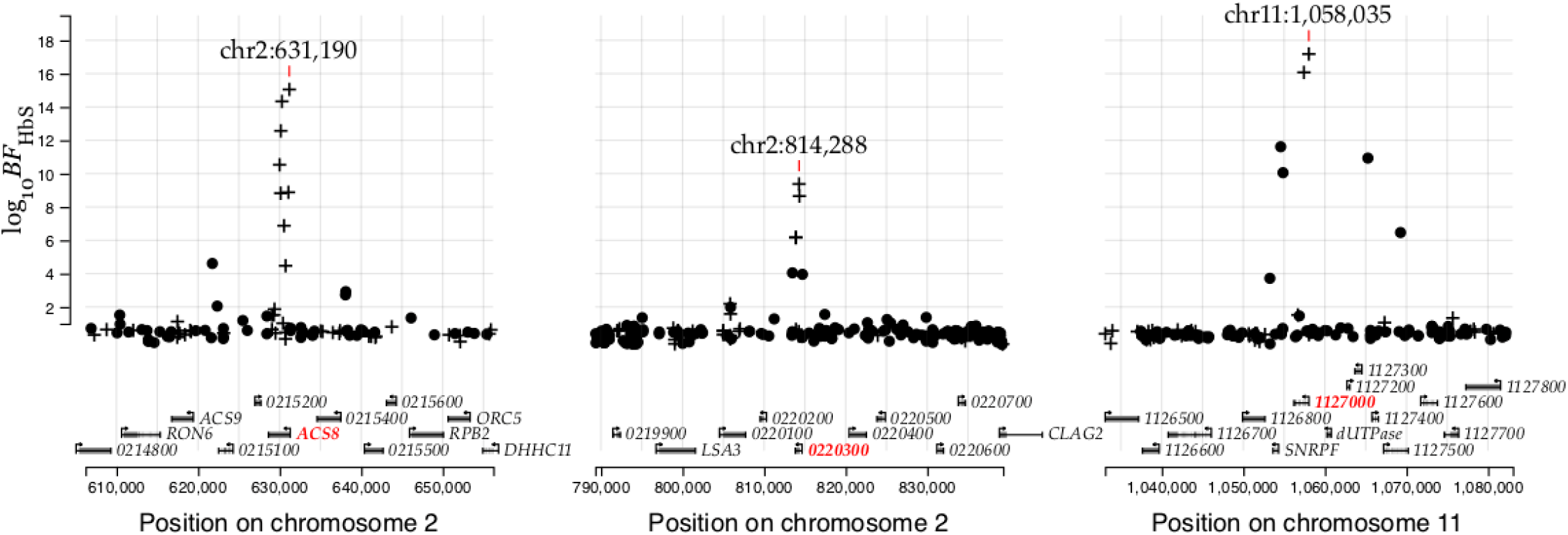
Evidence for association with HbS in three regions of the *Pf* genome. Points show evidence for association with HbS (log_10_ Bayes Factor for test in N=3,346 samples, y axis) for variants in the *Pfsa1*, *Pfsa2* and *Pfsa3* regions of the *Pf* genome (panels). Variants which alter protein coding sequence are denoted by plusses, while other variants are denoted by circles. Results are computed by logistic regression including an indicator of country as a covariate and assuming an additive model of association, with HbS genotypes based on imputation from genome-wide genotypes as previously described^5^; mixed and missing *Pf* genotype calls were excluded from the computation. A corresponding plot using directly-typed HbS genotypes can be found in **Supplementary Figure 5**. The variant with the strongest association in each region is annotated and the panels show regions of length 50kb centred at this variant. Below, regional genes are annotated, with gene symbols given where the gene has an ascribed name in the PlasmoDB annotation (after removing ‘PF3D7_’ from the name where relevant); the three genes containing the most-associated variants are shown in red.

We attempted to replicate this finding in a separate set of 825 samples in which the HbS genotypes have previously been assayed^2^ (**Supplementary Table 2**). The *Pfsa3* association replicated at nominal levels of evidence in the smaller Gambian sample (one-tailed *P* = 0.026), and all three loci replicated convincingly in the larger set of samples from Kenya (*P*< 0.001). Across the full dataset of 4,071 samples there is thus very strong evidence of association with HbS at all three loci (*BF*_HbS_ = 4.7×10^20^ for *Pfsa1*, 3.3×10^12^ for *Pfsa2*, and 2.5×10^24^ for *Pfsa3*; **Supplementary Figure 5**) with corresponding large effect size estimates (estimated odds ratio (OR) = 11.8 for *Pfsa1*+, 7.4 for *Pfsa2*+ and 21.7 for *Pfsa3*+). As described above, these estimates assume an additive relationship between HbS and the *Pf* genotype at each locus, but we also noted that genotype counts are most consistent with an overdominance effect (**Supplementary Figure 6**). We further examined the effect of adjusting for covariates including human and parasite principal components reflecting population structure, year of sampling, clinical type of severe malaria and technical features related to sequencing (**Supplementary Figure 7**). Inclusion of these covariates did not substantially affect results with one exception: we found that parasite principal components (PCs) computed across the whole *P.falciparum* genome in Kenya included components that correlated with the *Pfsa* loci, and including these PCs reduced the association signal. Altering the PCs by removing the *Pfsa* regions restored the association, indicating that this is not due to a general population structure effect that is reflected in genotypes across the *P.falciparum* genome, and we further discuss the reasons for this finding below. Taken together, these data appear to indicate genuine differences in the distribution of parasite genotypes between severe infections of HbS- and non-HbS genotype individuals.

The level of protection afforded by HbS can be estimated by comparing its frequency between severe malaria cases and population controls. As shown in **Figure 2**, the vast majority of children with HbS genotype in our data were infected with parasites that carry *Pfsa*+ alleles. Corresponding to this, our data show little evidence of a protective effect of HbS against severe malaria with parasites of *Pfsa1*+, *Pfsa2*+, *Pfsa3*+ genotype (estimated relative risk (*RR)* = 0.83, 95% CI = 0.53-1.30). In contrast, HbS is strongly associated with reduced risk of disease caused by parasites of *Pfsa1*-, *Pfsa2*-, *Pfsa3*- genotype (*RR =* 0.01, 95% CI = 0.007-0.03). These estimates should be interpreted with caution because they are based on just 49 cases of severe malaria that had an HbS genotype, because many of these samples were included in the initial discovery dataset, and because there is some variation evident between populations; however it can be concluded that the protective effect of HbS is dependent on parasite genotype at the *Pfsa* loci.

**Figure 2:**
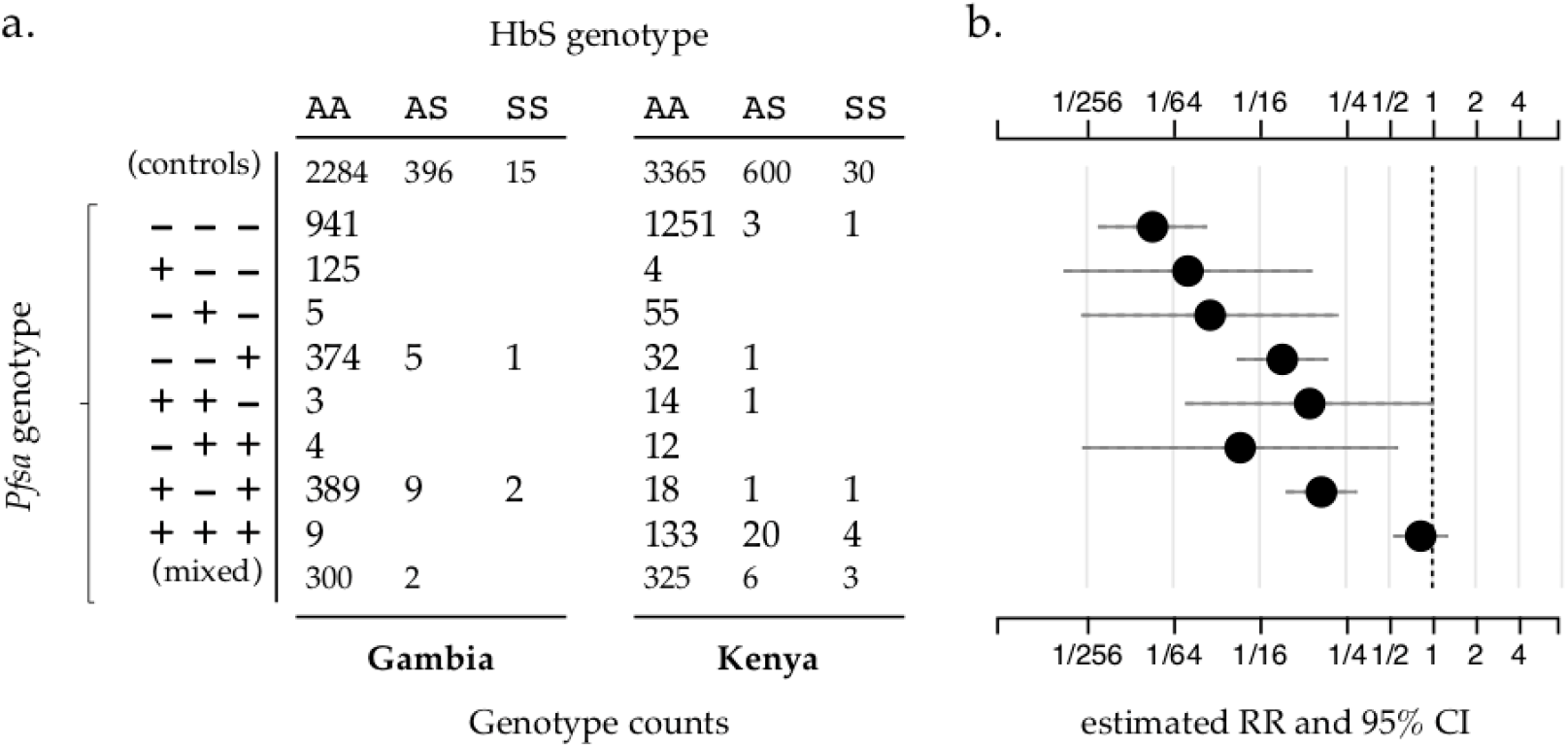
The estimated relative risk for HbS varies by *Pfsa* genotype. Panel a) shows the count of severe malaria cases from The Gambia and Kenya with given HbS genotype (columns; using *N =* 4,071 samples with directly-typed HbS genotype) and carrying the given alleles at the *Pfsa1*, 2, and 3 loci (rows). *Pfsa* alleles are indicated by + for the allele positively associated with HbS and - for the negatively associated allele at each locus. Samples with mixed *P.falciparum* genotype calls for at least one of the loci are shown in the last row and further detailed in **Supplementary Figure 8**. The first row indicates counts of HbS genotypes in population control samples from the same populations^5^. Panel b) shows the estimated relative risk of HbS on severe malaria with the given *Pfsa* genotypes (rows) using the data in panel a. Relative risks were estimated using a multinomial logistic regression model with controls as the baseline outcome and assuming complete dominance (i.e. that HbAS and HbSS genotypes have the same association with parasite genotype) as described in **Supplementary Methods**. An indicator of country was included as a covariate. To reduce overfitting we used Stan ^13^ to fit the model assuming a mild regularising Gaussian prior with mean zero and standard deviation of 2 on the log-odds scale (i.e. with 95% of mass between 1/50 and 50 on the relative risk scale) for each parameter, and between-parameter correlations set to 0.5. Solid horizontal lines denote the corresponding 95% credible intervals.

The *Pfsa1*+, *Pfsa2*+ and *Pfsa3*+ alleles had similar frequencies in Kenya (approximately 10-20%) whereas in Gambia *Pfsa2*+ had a much lower allele frequency than *Pfsa1*+ or *Pfsa3*+ (< 3% in all years studied, versus 25-60% for the *Pfsa1*+ or *Pfsa3*+ alleles; **Figure 3a and Supplementary Figure 9**). To explore the population genetic features of these loci in more detail, we analysed the MalariaGEN Pf6 open resource which gives *P. falciparum* genome variation data for 7,000 worldwide samples ^8^ (**Figure 3b**). This showed considerable variation in the frequency of these alleles across Africa, the maximum observed value being 61% for >*Pfsa3*+ in the Democratic Republic of Congo, and indicated that these alleles are rare outside Africa. Moreover, we found that within Africa, population frequencies of the *Pfsa*+ alleles are strongly correlated with the frequency of HbS (**Figure 3c**, estimated using data from the Malaria Atlas Project ^14^).

**Figure 3:**
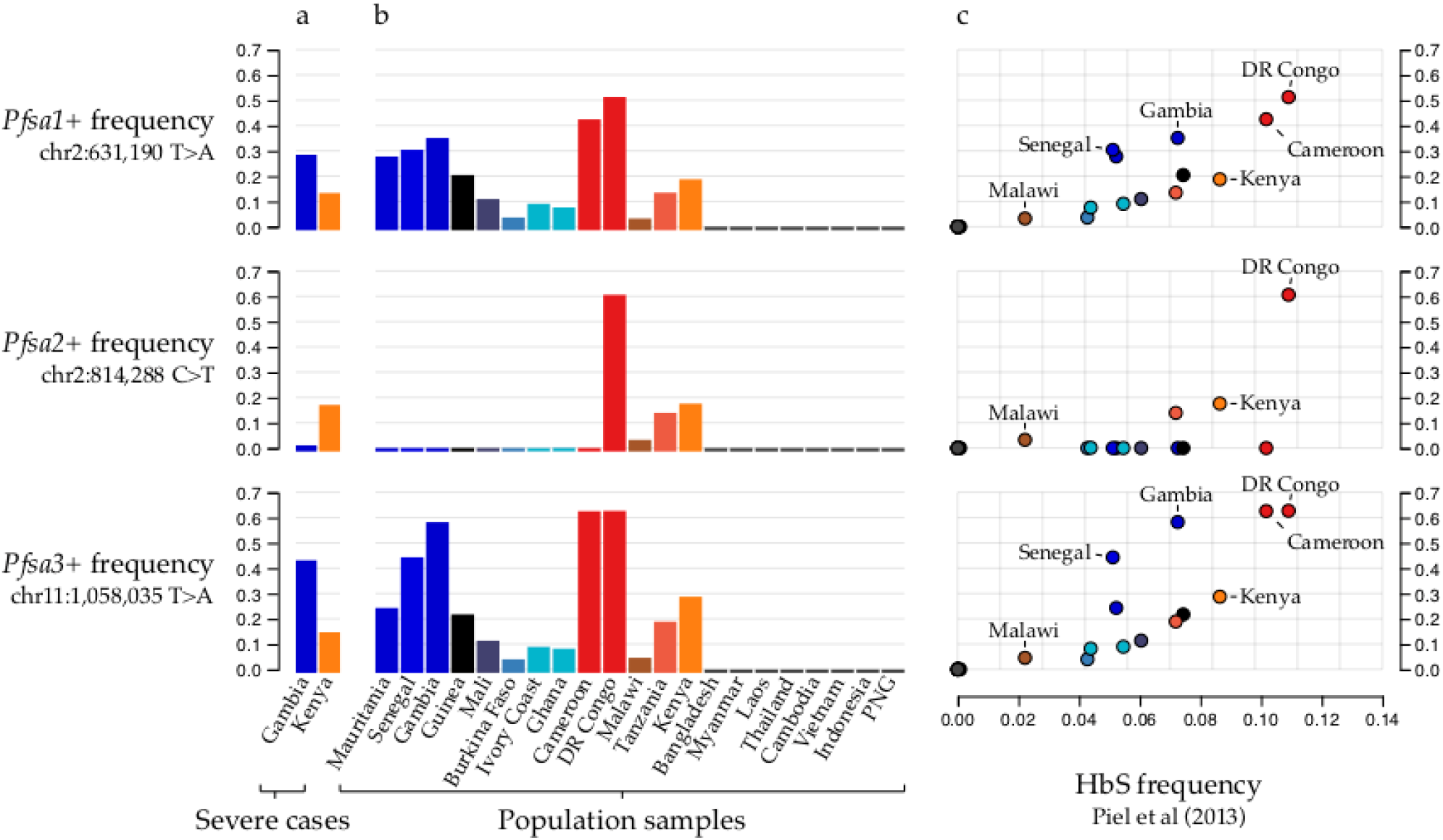
The relationship between *Pfsa* and HbS allele frequencies across populations. a) bars show the estimated frequency (y axis) of each *Pfsa*+ allele (rows) in severe malaria cases from each country (x axis and colours). Details of allele frequencies and sample counts across years of ascertainment can be found in **Supplementary Figure 9**. **b)** bars show the estimated frequency (y axis, as in panel a) of each *Pfsa*+ allele in worldwide populations from the MalariaGEN Pf6 resource, which contains samples collected in the period 2008-2015^8^. Only countries with at least 50 samples are shown (this excludes Columbia, Peru, Benin, Nigeria, Ethiopia, Madagascar, Uganda, and Bangladesh). **c)** Points show *Pfsa*+ allele frequency (y axis, as in panel a and b) against HbS allele frequency (x axis) in populations from MalariaGEN Pf6 (coloured as in panel b; selected populations are also labelled). HbS allele frequencies are computed from frequency estimates previously published by the Malaria Atlas Project^14^ by taking a weighted average over sampling sites within each country in MalariaGEN Pf6. All *Pfsa* allele frequencies were estimated after excluding mixed or missing genotype calls.

This analysis also revealed a further feature of the *Pfsa*+ alleles: although *Pfsa1* and *Pfsa2* are separated by 180kb, and the *Pfsa3* locus is on a different chromosome, they are in strong linkage disequilibrium (LD). This can be seen from the co-occurrence of these alleles in severe cases (**Figure 2),** and from the fact that they covary over time in our sample (**Supplementary Figure 9**) and geographically across populations (**Figure 3b**). To investigate this we computed LD metrics between the *Pfsa*+ alleles in each population (**Supplementary Table 3**) after excluding HbS-carrying individuals to avoid confounding with the association outlined above. *Pfsa1* + and *Pfsa2*+ were strongly correlated in Kenyan severe cases (r = 0.75) and *Pfsa1*+ and *Pfsa3*+ were strongly correlated in both populations (r = 0.80 in Kenya; r = 0.43 in severe cases from The Gambia). This high LD was also observed in multiple populations in MalariaGEN Pf6 (e.g. r = 0.20 between *Pfsa1*+ and *Pfsa3*+ in The Gambia; r = 0.71 in Kenya; r > 0.5 in all other African populations surveyed; **Supplementary Table 3**), showing that the LD is not purely an artifact of our severe malaria sample.

This observation of strong correlation between alleles at distant loci is unexpected, because the *P. falciparum* genome undergoes recombination in the mosquito vector and typically shows very low levels of LD in malaria endemic regions ^15–17^. To confirm that this is unusual, we compared LD between the *Pfsa* loci to the distribution computed from all common biallelic variants on different chromosomes (**Figure 4** and **Table 1**). In Kenyan samples, the *Pfsa* loci have the highest between-chromosome LD of any pair of variants in the genome. In Gambia, between-chromosome LD at these SNPs is also extreme, but another pair of extensive regions on chromosomes 6 and 7 also show strong LD (**Table 1**). These regions contain the chloroquine resistance-linked genes *pfCRT* and *pfAAT1* ^18,19^ and contain long stretches of DNA sharing identical by descent (IBD) consistent with positive selection of antimalarial-resistant haplotypes^20^. Moreoever, we noted that these signals are among a larger set of HbS-associated and drug-resistance loci that appear to have elevated between-chromosome LD in these data (**Supplementary Table 4**).

**Table 1:** Regions of highest correlation between *P.falciparum* chromosomes. Table shows all pairs of regions on different chromosomes containing pairs of SNPs with allele frequency at least 5% and squared correlation > 0.25 in each population. Region boundaries are defined to include all nearby pairs of correlated variants in either population with minor allele frequency >= 5% and r^2^ > 0.05, such that no other such pair of variants within 10kb of the given region boundaries is present. For each region in the pair, columns show the region boundaries, the lead variant, the region and/or gene containing the lead variant, the allele frequency, and the BF for association with HbS across populations. The rightmost columns give the sample size for the pairwise comparison after treating mixed genotype calls as missing, and the computed correlation. A longer list of regions showing between-chromosome LD can be found in **Supplementary Table 4**.

| Country | First region |  |  |  |  | Second region |  |  |  |  | Linkage disequilibrium |  |
| --- | --- | --- | --- | --- | --- | --- | --- | --- | --- | --- | --- | --- |
| | Region boundaries | Lead variant | Region / Gene | Allele frequency | $BF_{HbS}$ | Region boundaries | Lead variant | Region / Gene | Allele frequency | $BF_{HbS}$ | N | R |
| Gambia | chr6<br>1,174,040-1,293,328 | chr6<br>1,215,233<br>G > A | *PfAAT1* | 80% | 4.7 | chr7<br>361,356-481,853 | chr7<br>403,618<br>A > AT | *PfCRT* | 74% | 68 | 1,967 | 0.77 |
| Gambia | chr2<br>621,756-640,163 | chr2<br>631,092<br>T > C | *Pfsa1*<br>(*PfACS8*) | 28% | $5.8 \times 10^{12}$ | chr11<br>1,053,258-1,065,275 | chr11<br>1,057,437<br>T > C | *Pfsa3*<br>(1127000) | 46% | $8.7 \times 10^{21}$ | 1,881 | 0.44 |
| Kenya | chr2<br>621,756-631,190 | chr2<br>629,996<br>C > A | *Pfsa1*<br>(*PfACS8*) | 12% | $2.4 \times 10^{15}$ | chr11<br>1,053,258-1,069,278 | chr11<br>1,058,035<br>T > A | *Pfsa3*<br>(1127000) | 13% | $2.5 \times 10^{24}$ | 1,625 | 0.81 |
| Kenya | chr2<br>805,840-825,357 | chr2<br>814,329<br>A > G | *Pfsa2*<br>(0220300) | 16% | $2.2 \times 10^{12}$ | chr11<br>1,053,258-1,065,275 | chr11<br>1,057,437<br>T > C | *Pfsa3*<br>(1127000) | 14% | $8.7 \times 10^{21}$ | 1,557 | 0.67 |

**Figure 4:**
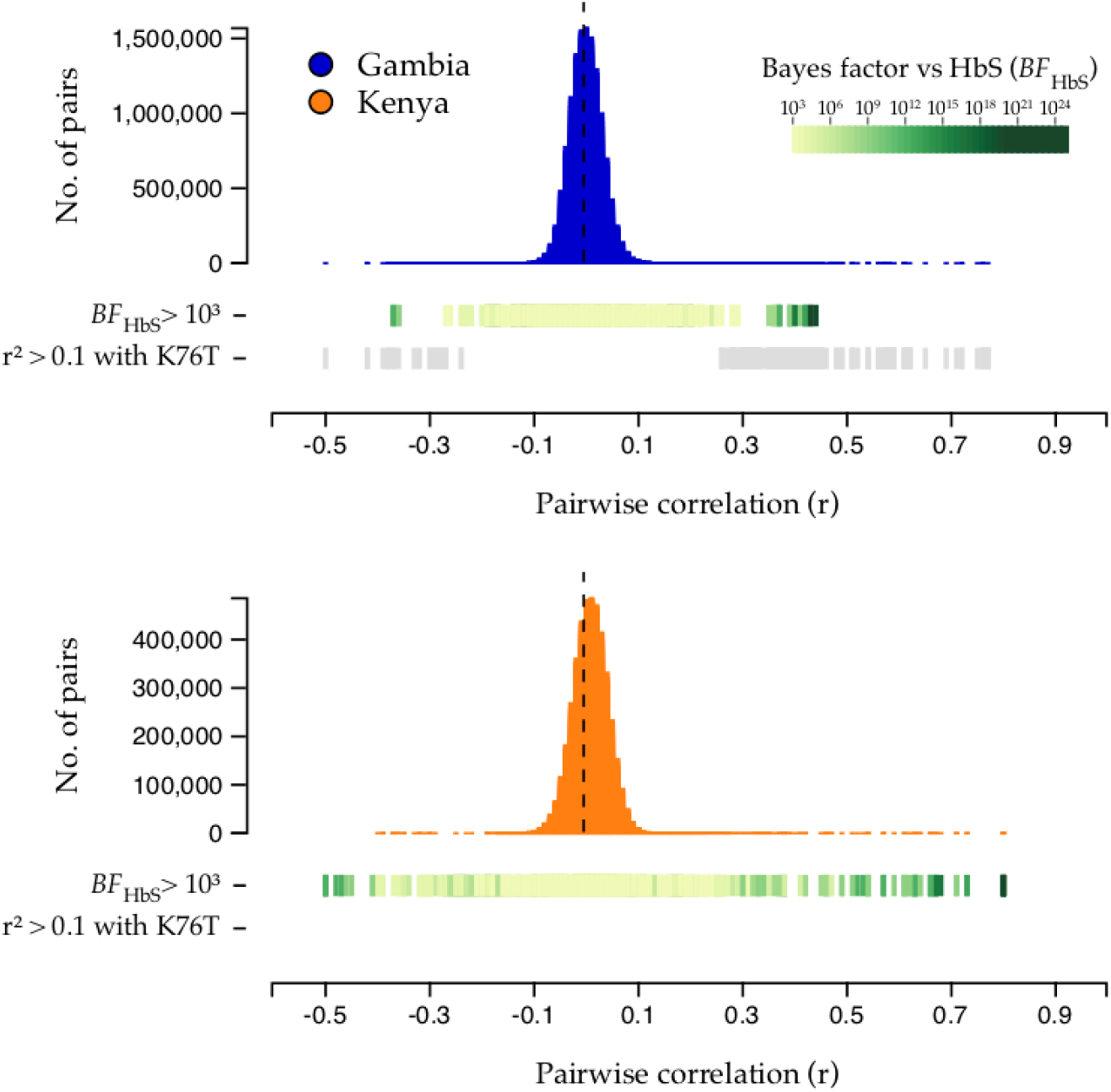
HbS-associated variants show extreme between-chromosome correlation in severe *P.falciparum* infections. Histograms show the distribution of genotype correlation (r) between variants on different *Pf* chromosomes in The Gambia (top panel; blue) and Kenya (bottom panel; orange). To avoid capturing effects of the HbS association, correlation values are computed after excluding HbS-carrying individuals. Correlation for each pair of variants is computed after excluding samples with mixed genotype calls, across all biallelic variants with estimated minor allele frequency at least 5% and at least 75% of samples having non-missing and non-mixed genotype call. Coloured bars indicate the evidence for association with HbS (*BF_HbS_*) for variants in each comparison (shown for variants with *BF_HbS_ >* 1,000; colour reflects the minimum *BF_HbS_* across the two variants in the pair as shown in the legend). Grey bars indicate variants having r^2^ > 0.1 with the *PfCRT* K76T mutation; as shown, no such variants were observed in Kenya.

Taking together these new findings with other population genetic evidence from multiple locations across Africa, including observations of frequency differentiation within and across *P.falciparum* populations ^17,21,22^ and other metrics at these loci indicative of selection ^20,23,24^, it appears likely that the allele frequencies and strong linkage disequilibrium between *Pfsa1*, *Pfsa2* and *Pfsa3* are maintained by natural selection. However, the mechanism for this is unclear. Given our findings, an obvious hypothesis is that the *Pfsa1*+, *Pfsa2*+ and *Pfsa3*+ alleles are positively selected in hosts with HbS, but since the frequency of HbS carriers is typically <20% ^2,14^ it is not clear whether this alone is a sufficient explanation to account for the high population frequencies or the strong LD observed in non-HbS carriers. Thus it remains entirely possible that there are other selective factors involved, such as epistatic interactions between these loci, or effects on fitness in the host or vector in addition to those observed here in relation to HbS.

The biological function of these parasite loci is a matter of considerable interest for future investigation. At the *Pfsa1* locus, the signal of association includes non-synonymous changes in the *PfACS8* gene, which encodes an acyl-CoA-synthetase ^25^. It belongs to a gene family that has expanded in the Laverania relative to other *Plasmodium* species^26^, and lies close to a paralog *PfACS9* on chromosome 2. The function of genes at the *Pfsa2* and *Pfsa3* loci are less well characterized. We analysed available genome assemblies of *P. falciparum* isolates^27^ and found evidence that *Pfsa3*+ is linked to a neighbouring copy number variant that includes duplication of the small nuclear ribonucleoprotein *SNRPF* (**Supplementary Figure 10**). Understanding the functional role of these loci could provide important clues into how HbS protects against malaria and help to distinguish between the various proposed mechanisms including: enhanced macrophage clearance of infected erythrocytes ^28^, inhibition of intraerythrocytic growth dependent on oxygen levels ^29^, altered cytoadherence of infected erythrocytes^30^ due to cytoskeleton remodelling ^31^ and immune-mediated mechanisms ^32^.

A fundamental question in the biology of host-parasite interactions is whether the genetic makeup of parasites within an infection is determined by the genotype of the host. While there is some previous evidence of this in malaria, e.g. allelic variants of the *PfCSP* gene have been associated with HLA type ^33^ and HbS has itself previously been associated with MSP-1 alleles ^34^, the present findings provide the clearest evidence to date of an interaction between genetic variants in the parasite and the host. Our central discovery is that, among African children with severe malaria, there is a strong association between HbS in the host and three loci in different regions of the parasite genome. Based on estimation of relative risk, HbS has no apparent protective effect against severe malaria in the presence of the *Pfsa1*+, *Pfsa2*+ and *Pfsa3*+ alleles. These alleles, which are much more common in Africa than elsewhere, are positively correlated with HbS allele frequencies across populations. However, they are found in substantial numbers of individuals without HbS as well, reaching up to 60% allele frequency in some populations. The *Pfsa1*, *Pfsa2* and *Pfsa3* loci also show remarkably high levels of long-range between-locus linkage disequilibrium relative to other loci in the *P. falciparum* genome, which is equally difficult to explain without postulating ongoing evolutionary selection. While it seems clear that HbS plays a key role in this selective process, there is a need for further population surveys (including asymptomatic and uncomplicated cases of malaria) to gain a more detailed understanding of the genetic interaction between HbS and these parasite loci, and how this affects the overall protective effect of HbS against severe malaria.

## Methods

### Ethics and consent

Sample collection and design of our case-control study^5^ was approved by Oxford University Tropical Research Ethics committee (OXTREC), Oxford, United Kingdom (OXTREC 020-006). Local approving bodies were the MRC/Gambia Government Ethics Committee (SCC 1029v2 and SCC670/630) and the KEMRI Research Ethics Committee (SCC1192).

### Building a combined dataset of human and *P.falciparum* genotypes in severe cases

We used Illumina sequencing to generate two datasets jointly reflecting human and *P.falciparum (Pf)* genetic variation, using a sample of severe malaria cases from The Gambia and Kenya for which human genotypes have previously been reported ^2,5^. A full description of our sequencing and data processing is given in **Supplementary Methods** and summarized in **Supplementary Figure 1**. In brief, following a process of sequence data quality control and merging across platforms, we generated i. a dataset of microarray and imputed human genotypes, and genome-wide *P.falciparum* genotypes, in 3,346 individuals previously identified as without close relationships^5^; and ii. a dataset of HbS genotypes directly typed on the Sequenom iPLEX Mass-Array platform (Agena Biosciences)^2^, and genome-wide *P.falciparum* genotypes, in 4,071 individuals without close relationships^5^. Parasite DNA was sequenced from whole DNA in samples with high parasitaemia, and using SWGA to amplify *Pf* DNA in all samples. *Pf* genotypes were called using an established pipeline^17^ based on GATK, which calls single nucleotide polymorphisms and short insertion/deletion variants relative to the Pf3D7 reference sequence. This pipeline deals with mixed infections by calling parasite variants as if the samples were diploid; in practice this means that variants with substantial numbers of reads covering reference and alternate alleles are called as heterozygous genotypes.

For the analyses presented in main text, we used the 3,346 samples with imputed human genotypes for our initial discovery analysis, and the 4,071 individuals with directly-typed HbS genotypes for all other analysis. The individuals in these two datasets substantially overlap (**Supplementary Figure 1**), but a subset of 825 individuals have directly-typed for HbS but were not in the discovery data and we used these for replication.

### Inference of genetic interaction from severe malaria cases

To describe our approach, we first consider a simplified model of infection in which parasites have a single definite (measurable) genotype, acquired at time of biting, that is relevant to disease outcome - i.e. we neglect any effects of within-host mutation, co- and super-infection at the relevant genetic variants. We consider the population of individuals who are susceptible to being been bitten by an infected mosquito, denoted *A*. A subset of infections go on to cause severe disease which we denote by *D*. Among individuals in *A* who are bitten and infected with a particular parasite type *I* = *y*, the association of a human allele *E* =*e* with disease outcome can be measured by the relative risk,

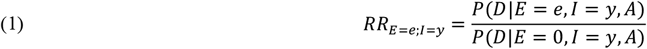

where we have used *E* = 0 to denote a chosen baseline human genotype against which risks are measured. If the strength of association further varies between parasite types then these relative risks will vary, such that the ratio of relative risks will differ from 1:

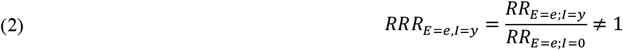

where we have used *I* = 0 to denote a chosen baseline parasite genotype. If the host genotype *e* confers protection against severe malaria, the ratio of relative risks will therefore capture variation in the level of protection compared between different parasite types.

Although expressed above in terms of a relative risk for human genotypes, rearrangement of terms in formula (2) can be equivalently expressed as a ratio of relative risks for a given parasite genotype compared between two human genotypes,

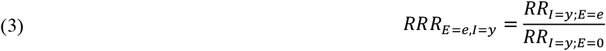

where *RR*_*I*=*j;E*=*e*_ is defined by analogy with (1). The ratio of relative risks is thus conceptually symmetric with respect to human and parasite alleles, and would equally well capture variation in the level of pathogenicity conferred by a particular parasite type compared between different human genotypes.

The odds ratio for specific human and parasite alleles computed in severe malaria cases is formally similar to the ratio of relative risks (2) but with the roles of the genotypes and *D* interchanged. Applying Bayes’ theorem to each term shows that in fact

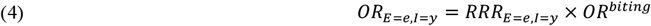

where *OR^biting^* is a term that reflects possible non-independence of human and parasite genotypes at the time of mosquito biting (**Supplementary Methods**). Thus, under this model, *OR*_*E*=*e*;*I*=*y*_ ≠1 implies either that host and parasite genotypes are not independent at time of biting, or that there is an interaction (on the risk scale; **Supplementary Methods**) between host and parasite genotypes in determining disease risk. The former possibility may be considered less plausible because it would seem to imply that relevant host and parasite genotypes can be detected by mosquitos prior to or during biting, but we stress that this cannot be tested formally without data on mosquito-borne parasites. A further discussion of these assumptions can be found in **Supplementary Methods**.

### Testing for genome-to-genome correlation

We developed a C++ program (HPTEST) to efficiently estimate the odds ratio (4) across multiple human and parasite variants. This program implements a logistic regression model in which genotypes from one file are included as the outcome variable and genotypes from a second file on the same samples are included as predictors. Measured covariates may also be included, and the model accounts for uncertainty in imputed predictor genotypes using the approach from SNPTEST^35^. The model is fit using a modified Newton-Raphson with line search method. For our main analysis we applied HPTEST with the parasite genotype as outcome and the host genotype as predictor, assuming an additive effect of the host genotype on the log-odds scale, and treating parasite genotype as a binary outcome (after excluding mixed and missing genotype calls.)

To mitigate effects of finite sample bias, we implemented regression regularised by a weakly informative log-F(2,2) prior distribution^36^ on the effect of the host allele (similar to a Gaussian distribution with standard deviation 1.87; **Supplementary Methods**). Covariate effects were assigned a log-F (0.08,0.08) prior, which has similar 95% coverage interval to a gaussian with zero mean and standard deviation of 40. We summarised the strength of evidence using a Bayes factor against the null model that the effect of the host allele is zero. A P-value can also be computed under an asymptotic approximation by comparing the maximum posterior estimate of effect size to its expected distribution under the null model (**Supplementary Methods**). For our main results we included only one covariate, an indicator of the country from which the case was ascertained (Gambia or Kenya); additional exploration of covariates is described below.

### Choice of genetic variants for testing

For our initial discovery analysis we concentrated on a set of 51,552 *Pf* variants that were observed in at least 25 individuals in our discovery set, after excluding any mixed or missing genotype calls. These comprised: 51,453 variants that were called as biallelic and passed quality filters (detailed in **Supplementary Methods**; including the requirement to lie in the core genome^37^); an additional 98 biallelic variants in the region of *PfEBL1* (which lies outside the core genome but otherwise appeared reliably callable); and an indicator of the *PfEBA175* ‘F’ segment, which we called based on sequence coverage as described in **Supplementary Methods and Supplementary Figure 11**. We included *PfEBL1* and *PfEBA175* variation because these genes encode known or putative receptors for *P.falciparum* during invasion of erythrocytes ^12^.

We concentrated on a set of human variants chosen as follows: we included the 94 autosomal variants from our previously reported list of variants with the most evidence for association with severe malaria^5^, which includes confirmed associations at *HBB*, *ABO*, *ATP2B4* and the glycophorin locus. We also included three glycophorin structural variants ^10^, and 132 HLA alleles (62 at 2-digit and 70 at 4-digit resolution) that were imputed with reasonable accuracy (determined as having minor allele frequency > 5% and IMPUTE info at least 0.8 in at least one of the two populations in our dataset). We tested these variants against all 51,552 *P.falciparum* variants described above. We also included all common, well-imputed human variants within 2kb of a gene determining a blood group antigen (defined as variants within 2kb of a gene in the HUGO Blood Group Antigen family^38^ and having a minor allele frequency of 5% and an IMPUTE info score of at least 0.8 in at least one of the two populations in our dataset; this includes 39 autosomal genes and 4,613 variants in total). We tested these against all variants lying within 2kb of *P.falciparum* genes previously identified as associated or involved in erythrocyte invasion ^11,12^ (60 genes, 1740 variants in total). In total we tested 19,830,288 distinct human-parasite variant pairs in the discovery dataset (**Supplementary Figure 4**).

### Definition of regions of pairwise association

We grouped all associated variant pairs (defined as pairs (*v,w)* having *BF(v,w)* > 100) into regions using an iterative algorithm as follows. For each associated pair *(v,w)*, we found the smallest enclosing regions (R_v_, R_w_) such that any other associated pair either lay with (R_v_, R_w_) or lay further than 10kb from (R_v_, R_w_) in the host or parasite genomes, repeating until all associated pairs were assigned to regions. For each association region pair, we then recorded the region boundaries and the lead variants (defined as the regional variant pair with the highest Bayes factor), and we identified genes intersecting the region and the gene nearest to the lead variants using the NCBI refGene^39^ and PlasmoDB v44^40^ gene annotations. Due to our testing a selected list of variant pairs as described above, in some cases these regions contain a single human or parasite variant. **Supplementary Table 1** summarises these regions for variant pairs with *BF* > 1,000.

### Frequentist interpretation of association test results

We compared association test P-values to the expectation under the null model of no association using a quantile-quantile plot, either before or after removing comparisons with HbS (**Supplementary Figure 3**; HbS is encoded by the ‘A’ allele at rs334, chr11:5,248,232 T -> A). A simple way to interpret individual points on the QQ-plot is to compare each P-value to its expected distribution under the relevant order statistic (depicted by the grey area in **Supplementary Figure 3**); for the lowest P-value this is similar to considering a Bonferroni correction.

### Bayesian interpretation of association test results

For each human variant *v*, we summarised the evidence that v is associated with variation in the parasite genome using an average Bayes factor computed across all the variants tested against v:

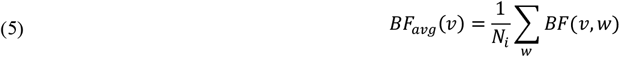

Here *BF*_*(v,w)*_ is the Bayes factor computed by HPTEST for the comparison between *v* and *w*, and the sum is over variants *w* in the parasite genome that were tested against *v*. Under the restrictive assumption that at most one parasite variant is associated with *w*, *BF*_*avg(v)*_ can be interpreted as a model-averaged Bayes factor reflecting the evidence for association; more generally *BF*_*avg*_ provides a pragmatic way to combines evidence across all tested variants. We similar define *BF*_*avg*_*(w)* for each parasite variant *w* averaged over all human variants tested against v. *BF*_*avg*_ is plotted for human and parasite variants in **Supplementary Figure 4**.

A direct interpretation of these Bayes factors requires assuming relevant prior odds. We illustrate this using a possible computation as follows. The 51,552 *Pf* variants represent around 20,000 1kb regions of the *Pf* genome, which might be thought of as approximately independent given LD decay rates 17. If we take the view that up to ten such regions might be associated with human genetic variants among those tested, this would dictate prior odds of around one in 2,000. With these odds, an average Bayes factor > 10,000 would be needed to indicate > 80% posterior odds of association. This calculation is illustrative; where specific information is available (for example, if a variant were known to affect a molecular interaction) this should be taken into account in the prior odds.

### Investigation of additional associations

In addition to the HbS-*Pfsa* associations, we also observed moderate evidence for association at a number of other variant pairs. These include associations between variation in the human gene *GCNT2* and *PfMSP4* with *BF* = 2.8×10^6^, and between HLA variation and multiple parasite variants with BF in the range 10^5^-10^6^ (**Supplementary Figure 4** and **Supplementary Table 1**). A fuller description of the context of these SNPs can be found in **Supplementary Methods**. Our interpretation is that the statistical evidence for these associations is not sufficiently strong on its own to make these signals compelling without additional evidence.

### Assessment of possible confounding factors

To assess whether the observed association between HbS and *P.falciparum* alleleles might be driven by confounding factors we conducted additional pairwise association tests as follows using HPTEST, based on directly-typed HbS genotypes and working seperately in the two populations. Results are shown in **Supplementary Figure 7**. First, we repeated the pairwise association test including only individuals overlapping the discovery dataset, and separately in the remaining set of 825 individuals. For discovery samples a set of population-specific principal components (PCs) reflecting human population structure were previously computed^5^ and we included these as covariates (including 20 PCs in total). Second, across all 4,071 individuals with directly-typed HbS data, we repeated tests including measured covariates as additional predictors. Specifically we considered: i. the age of individual at time of ascertainment (measured in years; range 0-12; treated as a categorical covariate), sex, reported ethnic group, and year of admission (range 1995-2010, treated as a categorical covariate); ii. technical covariates including an indicator of method of sequencing (SWGA or whole DNA), mean depth of coverage of the *Pf* genome, mean insert size computed from aligned reads, and percentage of mixed calls; and iii. an indicator of the clinical form of severe malaria which which the sample was ascertained (‘SM subtype’; either cerebral malaria, severe malarial anaemia, or other).

To assess the possibility that parasite population structure might impact results, we also included PCs computed in parasite populations as follows. Working in population separately, we started with the subset of biallelic SNPs with minor allele frequency at least 1% from among the 51,552 analysed variants (50,547 SNPs in Gambia and 48,821 SNPs in Kenya respectively). We thinned variants by iteratively picking variants at random from this list and excluding all others closer than 1kb (leaving 12,036 SNPs in Gambia and 11,902 SNPs in Kenya). We used QCTOOL to compute PCs using this list of SNPs. Several of the top PCs had elevated loadings from SNPs in specific genomic regions. This was especially noticeable in Kenya and included the widely-reported extensive regions of LD around the *AAT1* and *CRT* regions on chromosomes 6 and 7, and also the HbS-associated chromosome 2 and 11 loci. We therefore also considered separate sets of PCs computed after excluding SNPs in chromosomes 6 and 7 (leaving 9,933 and 9,812 SNPs respectively), after excluding chromosomes 2 and 11 (10,521 and 10,421 SNPs respectively) or after excluding 100kb regions centred on the lead HbS-associated SNPs (11,866 and 11,732 SNPs respectively). For each set of PCs, we repeated association tests including 20 PCs as fixed covariates.

For each subset of individuals, each HbS-associated variant and each set of covariates described above, we plotted the estimated effect size and 95% posterior interval, annotated with the total number of samples, the number carrying the non-reference allele at the given variant, and the number carrying heterozygous or homozygous HbS genotypes (**Supplementary Figure 7**). Corresponding genotype counts can be found in **Supplementary Figure 6.** To assess mixed genotypes calls, we also plotted the ratio of reads with reference and nonreference alleles at each site; this can be found in **Supplementary Figure 8**.

### Interpretation in terms of causal relationships

Observing *OR* ≠1 implies nonindependence between host and parasite genotypes in individuals with severe disease, but does not determine the mechanism by which this could occur. Assuming *OR*^biting^ = 1, we show in **Supplementary Methods** that *OR* = 1 is equivalent to the following multiplicative model of host and parasite genotypes on disease risk,

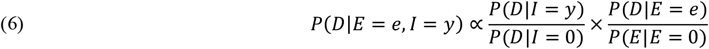

In general deviation from (6) could arise in several ways, including through within-host selection, interaction effects determining disease tolerance, as well as potential non-genotype-specific effects relating to disease diagnosis (similar to Berkson’s paradox ^41^). Our study provides only limited data to distinguish these possible mechanisms. For the HbS association described in main text, we note in **Supplementary Methods** that there is little evidence that the *Pfsa*+ variants are themselves associated with increased disease risk, and little evidence that the *Pfsa*+ variants associate with other host protective variants, suggesting that the observed interaction is specific to HbS.

### Comparison of severe cases to human population controls

Using D_y_ to denote severe disease caused by infection type y, the relative risk of the host genotype *E* = *e* on disease of type *y* can be written

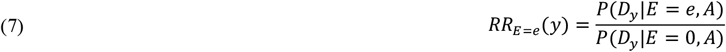

where *E* = 0 represents the baseline host genotype as above. Under the simplified infection model considered above, comparison with formula (1) relates this to the relative risk for host and parasite genotypes considered above,

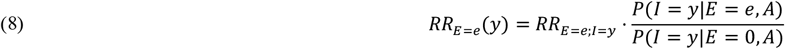

As in (4), the second term captures possible variation in infection rates for parasite type *y* between human genotypes, while the first term captures possible within-host effects. Direct comparison with (4) shows

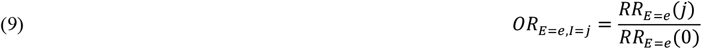

We show in **Supplementary Methods** that *RR_E=e_(y)* can be estimated using multinomial logistic regression comparing severe malaria cases to a sample of population controls, and we apply this approach in **Figure 2** to estimate *RR_E=e_(y)*, where *y* ranges over combined genotypes at the three *Pfsa* loci.

### Assessing sequencing performance in HbS-associated regions

We assessed sequencing performance at the chr2:631,190, chr2:814,288 and chr11:1,058,035 loci by computing counts of reads aligning to each position (“coverage”) and comparing this to the distribution of coverage across all biallelic sites in our dataset, treating each sample separately (**Supplementary Figure 11**). In general coverage at the three sites was high; we noted especially high coverage at chr2:814,288 in sWGA sequencing data (e.g. >90% of samples have coverage among the top 80% of that at biallelic variants genome-wide) but somewhat lower coverage in WGS samples at the chr11:1,058,035 locus. Variation in coverage between loci and samples is expected due to variation in DNA quantities, DNA amplification and sequencing processes, but we did not observe systeamtic differences in coverage between the different *Pfsa* genotypes at these loci. To further establish alignment accuracy, we also inspected alignment metrics and noted that across all analysis samples, over 99% of reads at each location carried either the reference or the identified non-reference allele, and over 99% of these reads had mapping quality at least 50 (representing confident read alignment). These results suggest sequencie reads provide generally accurate genotype calls at these sites.

### Assessing the distribution of between-chromosome LD

We developed a C++ program (LDBIRD) to efficiently compute LD between all pairs of *Pf* variants. LDBIRD computes the frequency of each variant, and computes the correlation between genotypes at each pair of variants with sufficiently high frequency. It then generates a histogram of correlation values and reports pairs of variants with squared correlation above a specified level. We applied LDBIRD separately to *Pf* data from Gambian and Kenyan severe malaria cases. We restricted attention to comparisons between biallelic variants that had frequency at least 5% in the given population and with at least 75% of samples having non-missing genotypes at both variants in the pair, after treating mixed genotype calls as missing, and output all pairs with r^2^ at least 0.01 for further consideration. To avoid confounding of LD by the HbS association signal, we also repeated this analysis after excluding individuals that carry the HbS allele (with the latter results presented in **Figure 4 and Supplementary Table 2**).

To summarise between-chromosome LD results we grouped signals into regions as follows. First, we observed that most variant pairs have |r| < 0.15 and hence r^2^ > 0.05 is typically a substantially outlying degree of inter-chromosomal LD (Figure 4). We therefore focussed on variant pairs (v1,v2) with r^2^ > 0.05. To each such pair (v1,v2) we assigned a pair of LD regions (R1,R2) with the property that R1 and R2 capture all other nearby variants with high r^2^. Specifically, R1 and R2 are defined as the smallest regions containing v1 and v2 respectively, such that for every other pair of variants (w1,w2) on the same chromosomes with r^2^>0.05,

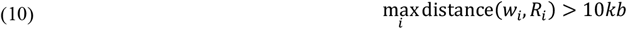

To compute R1 and R2, we implemented an iterative algorithm that successively expands the initial pair until no additional nearby pairs with high r^2^ can be found.

For each LD region pair we recorded the region boundaries and the most-correlated pair of variants. For Table 1 we list the region pairs with r^2^ > 0.25, reporting the superset of the region boundaries defined in the Gambian and Kenyan data where applicable. A full list of region pairs with r^2^ > 0.05 is given in **Supplementary Table 3**.

### Assessing the structure of *Pfsa* regions in available genome assemblies

We extracted 101bp and 1001bp flanking sequence centred at the chr2:631,190, chr2:814,288 and chr11:1,058,035 loci from the Pf3D7 reference sequence. We then used minimap2 ^42^ to align these sequences to a previously generated set of genome assemblies from *P.falciparum* isolates and laboratory strains ^27^ (**Supplementary Table 4**), allowing for multiple possible mapping locations. Each flanking sequence aligned to a single location on the corresponding chromosome in all included genomes, with the exception that sequence flanking the chromosome 11 locus aligned to two locations in the ML01 sample. This sample was excluded from previous analysis^27^ as it represents a multiple infection; we comment further on this below.

To further inspect sequence identity, we used MAFFT to generate a multiple sequence alignment (MSA) corresponding to the 1001bp sequence centred at each locus. Four isolates (GA01 from The Gabon, SN01 from Senegal, Congo CD01 and ML01 from Mali) carry the non-reference ‘A’ allele at the chr11:1,058,035 SNP; two of these (GA01 and CD01) also carry the non-reference allele at the chr2:631,190 SNP and one (CD01) carries the non-reference allele at all three SNPs. However, expansion of alignments to include a 10,001bp segment indicated that these four samples also carry a structural rearrangement at the chr11 locus. Specifically, GA01, SN01, CD01 and ML01 genomes include a ~1kb insertion present approximately 900bp to the right of chr11:1,058,035, and also a ~400bp deletion approximately 2400bp to the left of chr11:1,058,035. To investigate this, we generated kmer sharing ‘dot’ plots for k=50 across the region (**Supplementary Figure 10**), revealing a complex rearrangement carrying both deleted and duplicated segments. The duplicated sequence includes a segment (approx. coordinates 1,054,000-1,055,000 in Pf3D7) that contains the gene *SNRPF* (‘small nuclear ribonucleoprotein F, putative’) in the Pf3D7 reference. Inspection of breakpoints did not reveal any other predicted gene copy number changes in this region, including for *Pf3D7_1127000*.

As noted above, the chromosome 11 region aligns to a second contig in ML01 (contig chr0_142, **Supplementary Table 4**). This contig appears to have a different tandem duplication of a ~4kb segment lying to the right of the associated SNP (approximately corresponding to the range 11:1,060,100 – 1,064,000 in Pf3D7; Supplementary Figure 8). This segment contains a number of genes including dUTPase, which has been under investigation as a potential drug target^43^. We interpret this second contig as arising due to the multiple infection in this sample^27^, and given challenges inherent in genome assembly of mixed samples it is unclear whether this duplication represents an assembly artefact or a second genuine regional structural variant.

## Data Availability

A full list of data generated by this study and relevant accessions can be found at http://www.malariagen.net/resource/32.

## Code Availability

Source code for HPTEST and LDBIRD is available at https://code.enkre.net/qctool under an open-source license.

## Author contributions

Conceptualization: G.B., E.M.L., T.N.W., K.A.R., D.P.K.; Data Curation: G.B., E.M.L., T.N., M.J., C.M.N., R.D.P., R.A., K.A.R.; Formal Analysis: G.B., E.M.L., K.A.R.; Funding Acquisition: D.P.K; Investigation: C.H., A.E.J., K.R., E.D., K.A.R.; Methodology: G.B., K.A.R., D.P.K; Project Administration: S.M.G., E.D., K.A.R., D.P.K.; Resources: S.M.G., E.D., J.S., C.V.A., R.A., R.D.P., M.J., F.S-J., K.A.B., G.S., C.M.N., A.W.M., N.P., C.H., A.E.J., K.R., E.D., K.A.R.; Software and visualisation: G.B.; Supervision: D.J.C., U.d’A., K.M., T.N.W., S.M.G., K.A.R., D.P.K; Writing: G.B., E.M.L., T.N.W., K.A.R., D.P.K. in collaboration with all authors.

## Acknowledgements

We thank the patients and staff of Kilifi County Hospital and the KEMRI-Wellcome Trust Research Programme, Kilifi for their help with this study, and members of the Human Genetics Group in Kilifi for help with sample collection and processing. We thank the patients and staff at the Paediatric Department of the Royal Victoria Hospital in Banjul, Gambia for their help with the study. The human genetic data used in this study has previously been reported by the Malaria Genomic Epidemiology Network, and we thank all our colleagues who contributed to this previous work as part of MalariaGEN Consortial Project 1. A full list of consortium members is provided at https://www.malariagen.net/projects/consortial-project-1/malariagen-consortium-members. The MalariaGEN Pf6 open resource^17^ was generated through the Malaria Genomic Epidemiology Network *Plasmodium falciparum* Community Project (https://www.malariagen.net/resource/26).

The Malaria Genomic Epidemiology Network study of severe malaria was supported by Wellcome (https://wellcome.ac.uk/) (WT077383/Z/05/Z [MalariaGEN]) and the Bill & Melinda Gates Foundation ( https://www.gatesfoundation.org/) through the Foundations of the National Institutes of Health (https://fnih.org/) (566 [MalariaGEN]) as part of the Grand Challenges in Global Health Initiative. The Resource Centre for Genomic Epidemiology of Malaria is supported by Wellcome (090770/Z/09/Z; 204911/Z/16/Z [MalariaGEN]). This research was supported by the Medical Research Council (https://mrc.ukri.org/) (G0600718; G0600230; MR/M006212/1 [MalariaGEN]). Wellcome also provides core awards to the Wellcome Centre for Human Genetics (203141/Z/16/Z [WCHG]) and the Wellcome Sanger Institute (206194 [WSI]). Genome sequencing was carried out at the Wellcome Sanger Institute and we thank the staff of the Wellcome Sanger Institute Sample Logistics, Sequencing, and Informatics facilities for their contribution. TNW is supported through a Senior Fellowship from Wellcome (202800/Z/16/Z). This paper is published with permission from the Director of the Kenya Medical Research Institute (KEMRI). This research was funded in whole or in part by Wellcome as detailed above. For the purpose of Open Access, the author has applied a CC-BY public copyright licence to any author accepted manuscript version arising from this submission. The funders had no role in study design, data collection and analysis, decision to publish, or preparation of the manuscript.

## Notes

### Competing Interest Statement

The authors have declared no competing interest.

