## Supplementary Figures 1-12 for "The protective effect of sickle cell haemoglobin against severe malaria depends on parasite genotype"

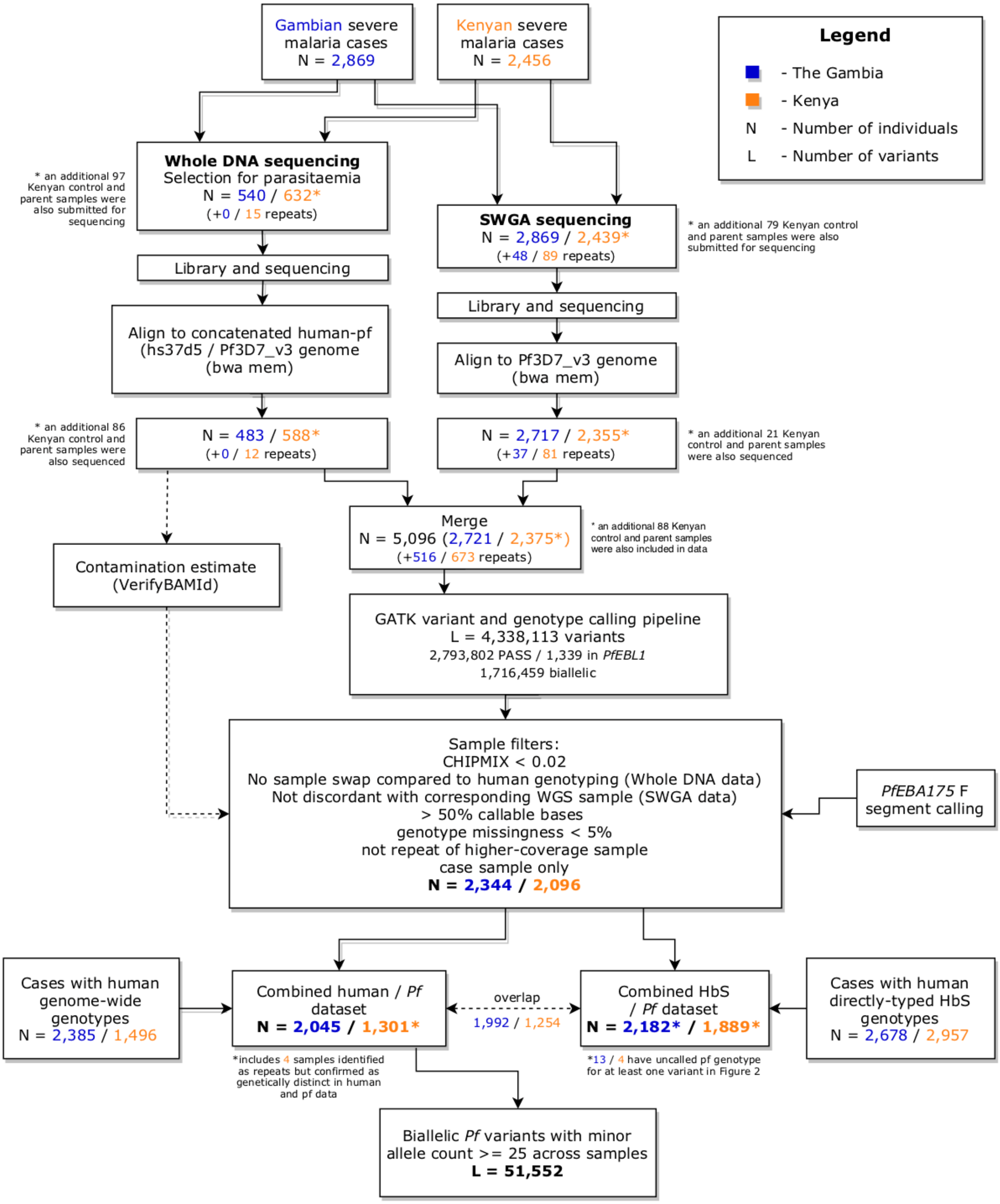

**Supplementary Figure 1. Flowchart showing generation and processing of *P.falciparum* (*Pf*) sequence data from 5,096 severe malaria cases.**

Flowchart shows sample processing from initial selection for whole DNA and Selective Whole genome Amplification (SWGA) pipelines (top) to the curated analysis datasets (bottom). Numbers in each box show counts of severe cases in The Gambia (blue) and Kenya (orange) with the number of individuals sequenced multiple times indicated in brackets. Following curation and QC of data (large box), available *Pf* data was intersected with two existing human genotype datasets to for the analyses described in main text. The combined pf/human imputed dataset has 3,346 samples and the combined pf/human direct typing dataset contains 4,071 individuals. These two datasets have substantial overlap; 825 individuals were represented in the directly-typed data but not the imputed data and were used for replication.

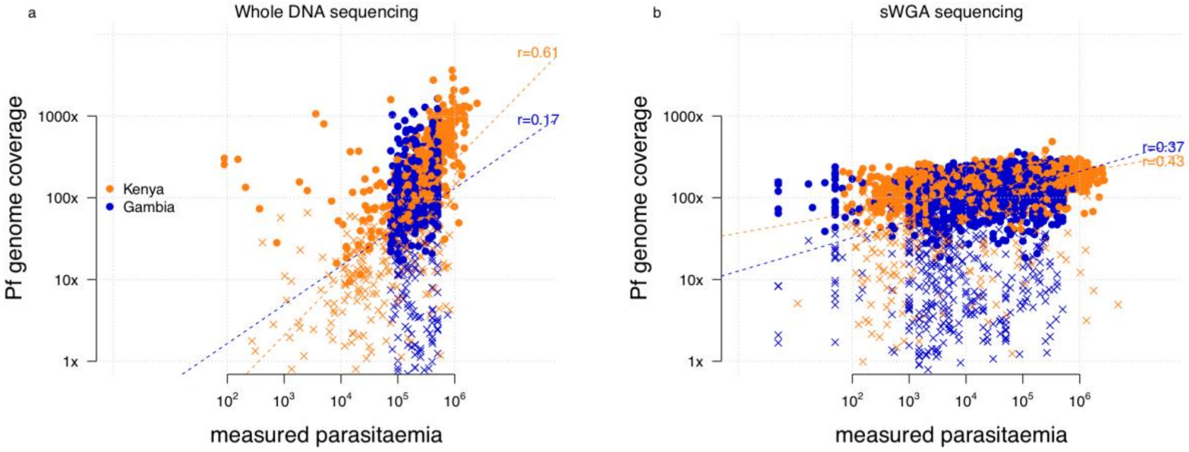

**Supplementary Figure 2. *Pf* genome coverage compared to measured parasitaemia in Whole DNA and sWGA sequencing pipelines.**

For each sample (points) sequenced using the Whole DNA pipeline (panel a) or sWGA pipeline (panel b), figure shows the average per-base coverage of reads aligned to the Pf3D7 genome (y axis) against the *P.falciparum* parasitaemia (parasitsed RBCs / ul blood) measured using blood slide at the time of ascertainment in each country (colours). Crosses denote samples that failed QC metrics (detailed in **Supplementary Methods** and **Supplementary Figure 1**) and were excluded from our analysis dataset. Lines show linear regression fit of $\log_{10}$(coverage) against $\log_{10}$(parasitaemia), with labels indicating the estimated correlation.

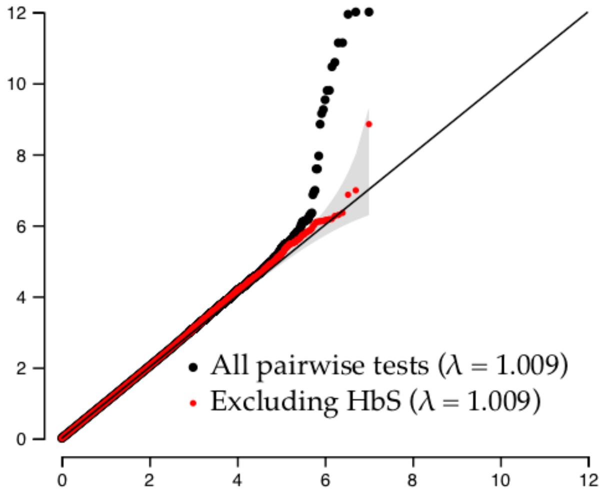

**Supplementary Figure 3. Quantile-quantile plot for test of association between pairs of human and P.falciparum alleles**

Plot shows observed -log10 P-value (y axis) against expection under the null model of no association (x axis) for tests of association between human and pf alleles. Tests are conducted using logistic regression across 3,346 samples, with the imputed human genotype as predictor and the parasite genotype as outcome. An indicator of country (Gambia or Kenya) was included as a covariate. Black (respectively red) points reflect quantile-quantile plot for all tests (black) or after excluding comparisons with HbS (red points), with the corresponding median lambda values shown in the legend. Grey area depicts the 99% confidence interval for the observed value, computed pointwise for each black point using the order statistics for a uniform distribution. Only comparisons where the minor allele count of the human variant for samples carrying either *Pf* genotype is at least 20 are shown (computed in expectation across the imputed genotype distribution, where relevant).

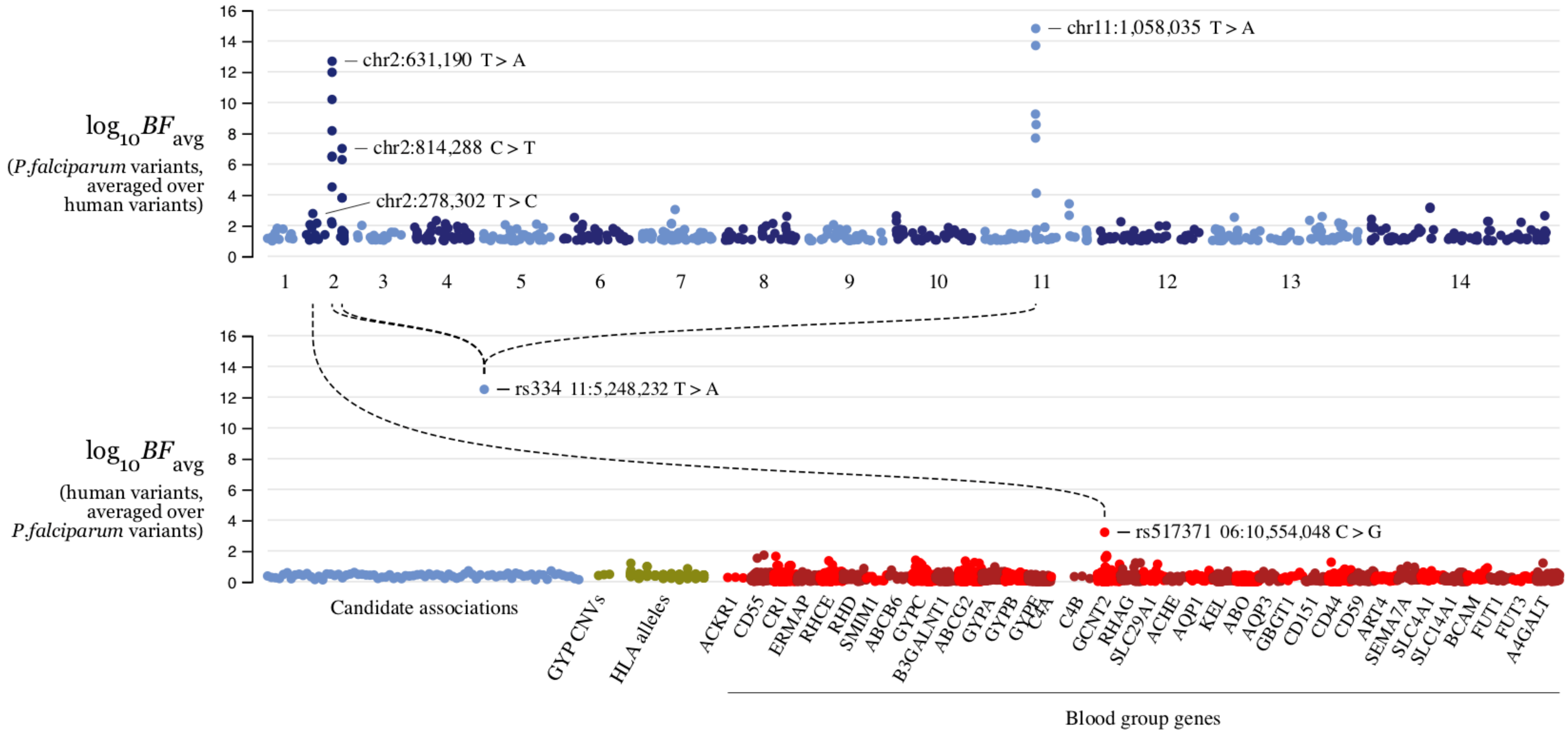

**Supplementary Figure 4. Summary of association between parasite variants and human variants.**

Figure shows model-averaged BF ($BF_{avg}$; Methods) computed for each parasite variant (top panel) and for each human variant (bottom panel) tested. For each variant, $BF_{avg}$ is computed by averaging the per-test BF across all variants in the other organism against which the variant was tested, as described in Methods. Pf variants are organised by location in the Pf genome (top panel). Human variants are organised by category; we considered previously established or suggestive associations [1], common Glycophorin copy number variants [2], imputed HLA alleles [1]; we also considered tests between variants within 2kb of red blood group genes and Pf variants within 2kb of previously identified proteins important for blood stage parastes[3,4] as described in Methods. Dashed lines denote individual associations with $BF > 1{,}000{,}000$; the lead variants in the relevant regions are annotated.

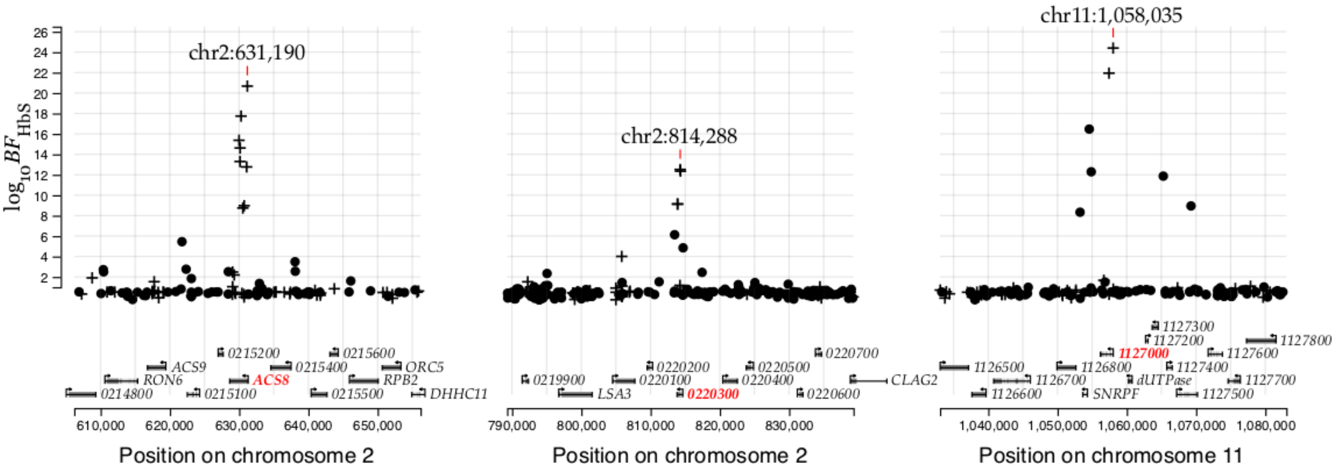

**Supplementary Figure 5. Evidence for association with HbS in three regions of the *Pf* genome using directly-typed HbS genotypes.**

Points show evidence for association with HbS ($log_{10}$ Bayes Factor for test in N=4,071 samples, y axis) based on direct typing of HbS for variants in the *Pfsa1*, *Pfsa2* and *Pfsa3* regions of the *Pf* genome (panels). Variants which alter protein coding sequence are denoted by plusses, while other variants are denoted by circles. Results are computed by logistic regression including an indicator of country as a covariate and assuming an additive model of association; missing and mixed Pf genotype calls were excluded. A corresponding plot using imputed HbS genotypes can be found in Figure 1. The variant with the strongest association in each region is annotated and the panels show regions of length 50kb centred at this variant. Below, regional genes are annotated, with gene symbols given where the gene has an ascribed name in the PlasmoDB annotation (after removing 'PF3D7_' from the name where relevant); the three genes containing the most-associated variants are shown in red.

### chr2:631,190 T/A

| | HbS genotype | | | | | |
|---|---|---|---|---|---|---|
| | Gambia | | | Kenya | | |
| | A/A | A/T | T/T | A/A | A/T | T/T |
| T | 1447 | 5 | 1 | 1504 | 4 | 1 |
| T/A | 155 | 1 | | 144 | 3 | 1 |
| A | 560 | 10 | 2 | 200 | 25 | 7 |
| (controls) | 2284 | 396 | 15 | 3365 | 600 | 30 |

### chr2:814,288 C/T

| | HbS genotype | | | | | |
|---|---|---|---|---|---|---|
| | Gambia | | | Kenya | | |
| | A/A | A/T | T/T | A/A | A/T | T/T |
| C | 2124 | 15 | 3 | 1357 | 6 | 2 |
| C/T | 16 | 1 | | 244 | 2 | 1 |
| T | 23 | | | 247 | 24 | 6 |
| (controls) | 2284 | 396 | 15 | 3365 | 600 | 30 |

### chr11:1,058,035 T/A

| | HbS genotype | | | | | |
|---|---|---|---|---|---|---|
| | Gambia | | | Kenya | | |
| | A/A | A/T | T/T | A/A | A/T | T/T |
| T | 1111 | | | 1463 | 5 | 1 |
| T/A | 212 | | | 161 | 3 | 2 |
| A | 828 | 16 | 3 | 220 | 24 | 6 |
| (controls) | 2284 | 396 | 15 | 3365 | 600 | 30 |

**Supplementary Figure 6. Counts of combined human and pf genotypes observed in severe malaria cases, by variant**

Panels show counts of severe malaria cases in The Gambia and Kenya with the given HbS genotype (columns) and given infection genotype (rows) at each of the lead variants in the *Pfsa* regions (panels). Counts are computed in the directly-typed HbS genotype dataset.

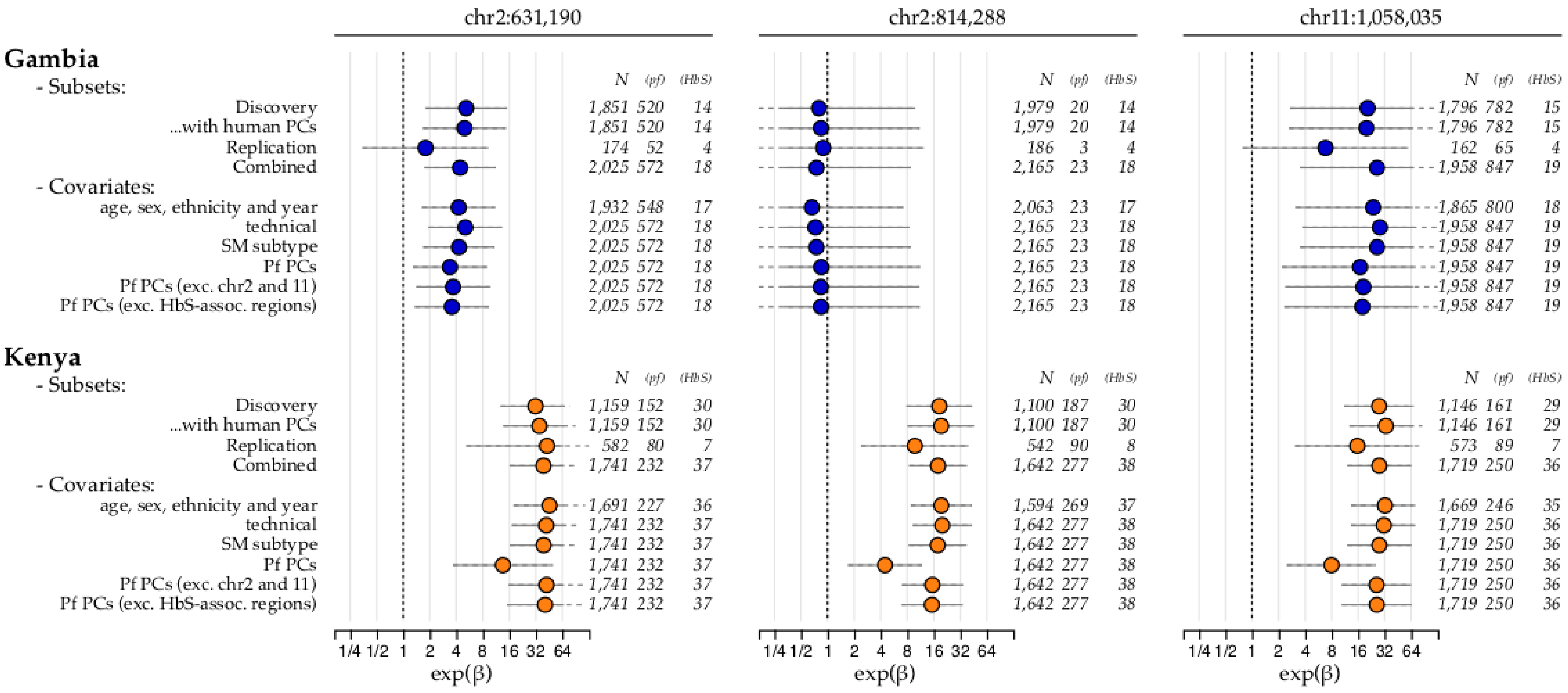

**Supplementary Figure 7. Odds ratios for association of HbS with the *Pfsa* variants in severe malaria cases.**

Plot shows parameter estimates (points) and 95% posterior credible intervals (horizontal line segments) for the association of HbS with *Pf* genotype at each of the three *Pfsa* lead variants (columns), using several combinations of sample subsets and covariates (rows) in The Gambia and Kenya. Estimates are computed separately for each SNP using logistic regression with the given covariates included as fixed-effect terms, and are based on directly-typed HbS genotypes assuming a dominance model of HbS on *Pf* genotype. Samples with mixed *Pf* genotype calls are excluded from the regression. All estimates are made using a weakly-informative log-F*(2,2)* prior (**Supplementary Methods**); a diffuse log-F*(0.08,0.08)* prior is also applied to covariate effects. Row names are as follows: "Discovery": samples with human genome-wide imputed data that were included in our initial scan (**Supplementary Figure 4 and Figure 1**); "Replication": the 825 additional samples that are not closely related to discovery samples (as determined previously[1]); "Combined": all samples with direct typing (as in **Figure 2 and Supplementary Figure 5**); "technical": indicators of sequencing performance including indicator of SWGA or whole DNA sequencing method for the sample, sequence read depth, insert size, and proportion of mixed genotype calls; "SM subtype": indicator of clinical presentation (cerebral malaria, severe malarial anaemia or other severe malaria) the individual was ascertained with; "Pf PCs": principal components (PCs) computed using all called biallelic SNPs having minor allele frequency at least 1% in each population and thinned to exclude variants closer than 1kb; additional rows are shown for PCs computed after excluding SNPs in chromosomes 2 and 11, or from the three regions of association shown in **Figure 1** plus a 25kb margin. Numbers to the right of each estimate show the total regression sample size, the number of samples having the non-reference allele at the given *Pf* SNP, and the number heterozygous or homozygous for HbS.

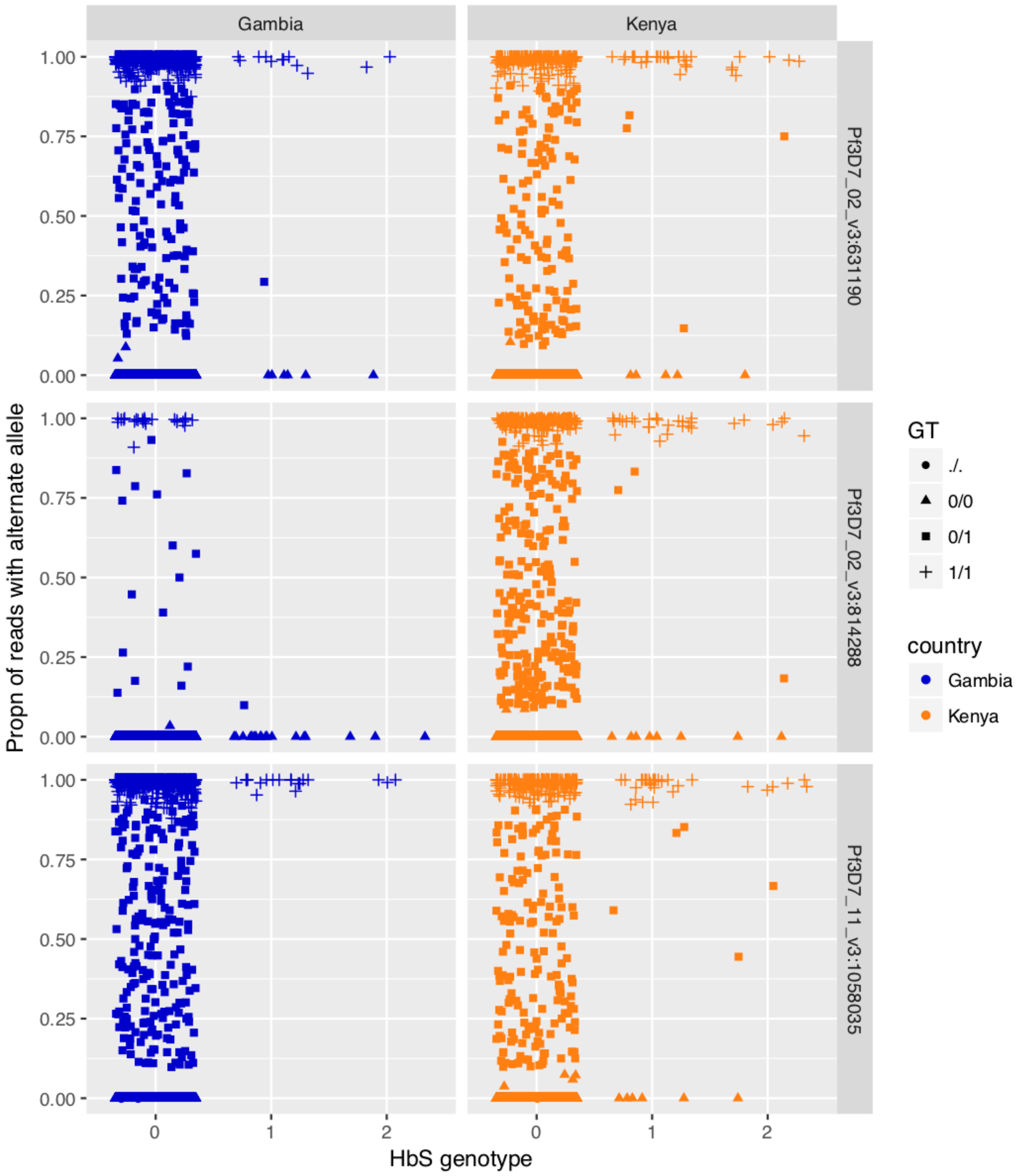

**Supplementary Figure 8. Allele read ratio versus HbS genotype at the three HbS-associated loci**

For each sample (points) and each of the three HbS-associated loci (rows), the figure shows the proportion of sequencing reads that carry the nonreference allele (y axis). Points are separated by country (columns) and HbS genotype (x axis); the x axis values are jittered to visually separate. The called *Pf* genotype of each sample is indicated by the shape, with mixed calls indicated by squares.

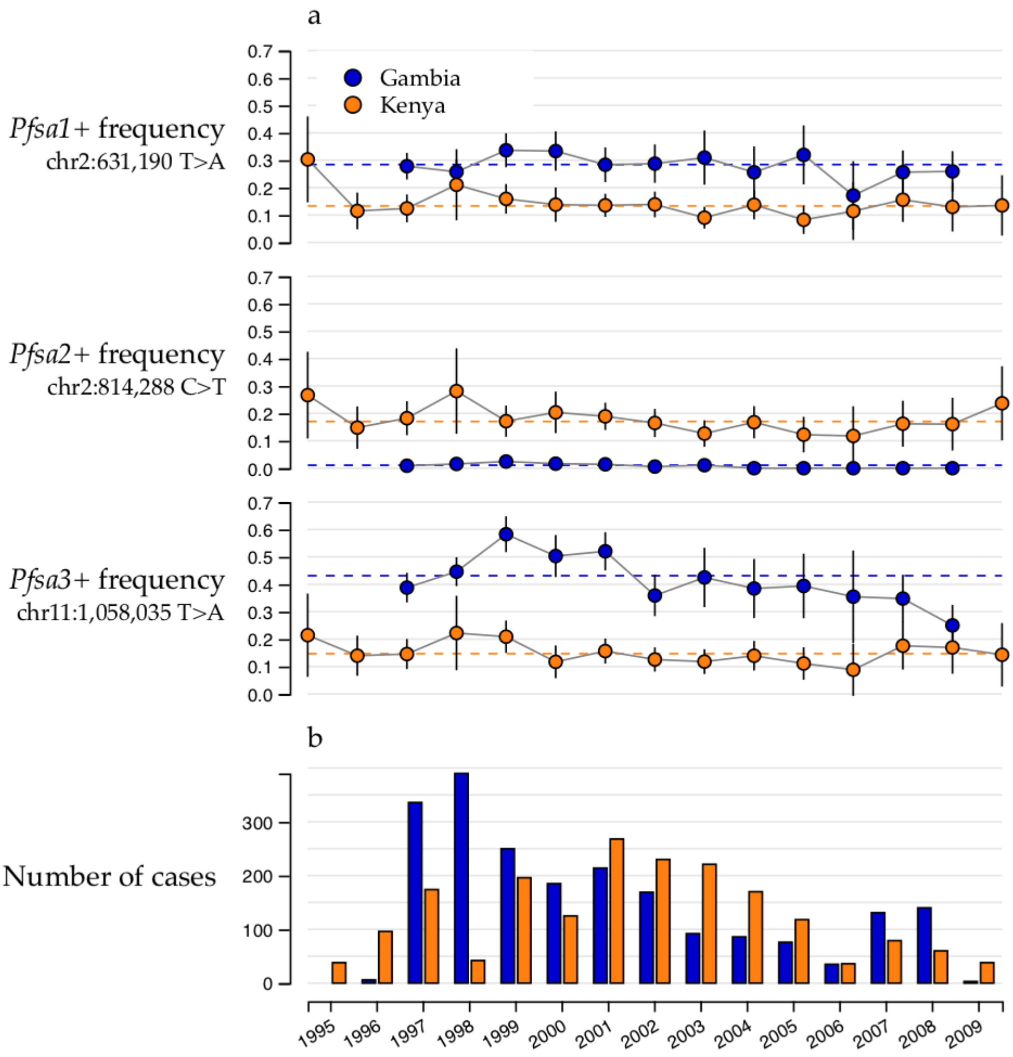

**Supplementary Figure 9. *Pfsa*+ allele frequency and sample size by year of ascertainment**

**a)** points show the sample allele frequency (y axis) for each Pfsa variant (rows) in severe malaria cases by year of ascertainment (x axis) and country (colour). Vertical line segments show the 95% confidence interval corresponding to each estimate. Horizontal dashed lines show the overall estimate across all years in our data, as in **Figure 3a**. **b)** Bars show the total number of severe case samples in our dataset (y axis) in each country (colour) by year of ascertainment (x axis).

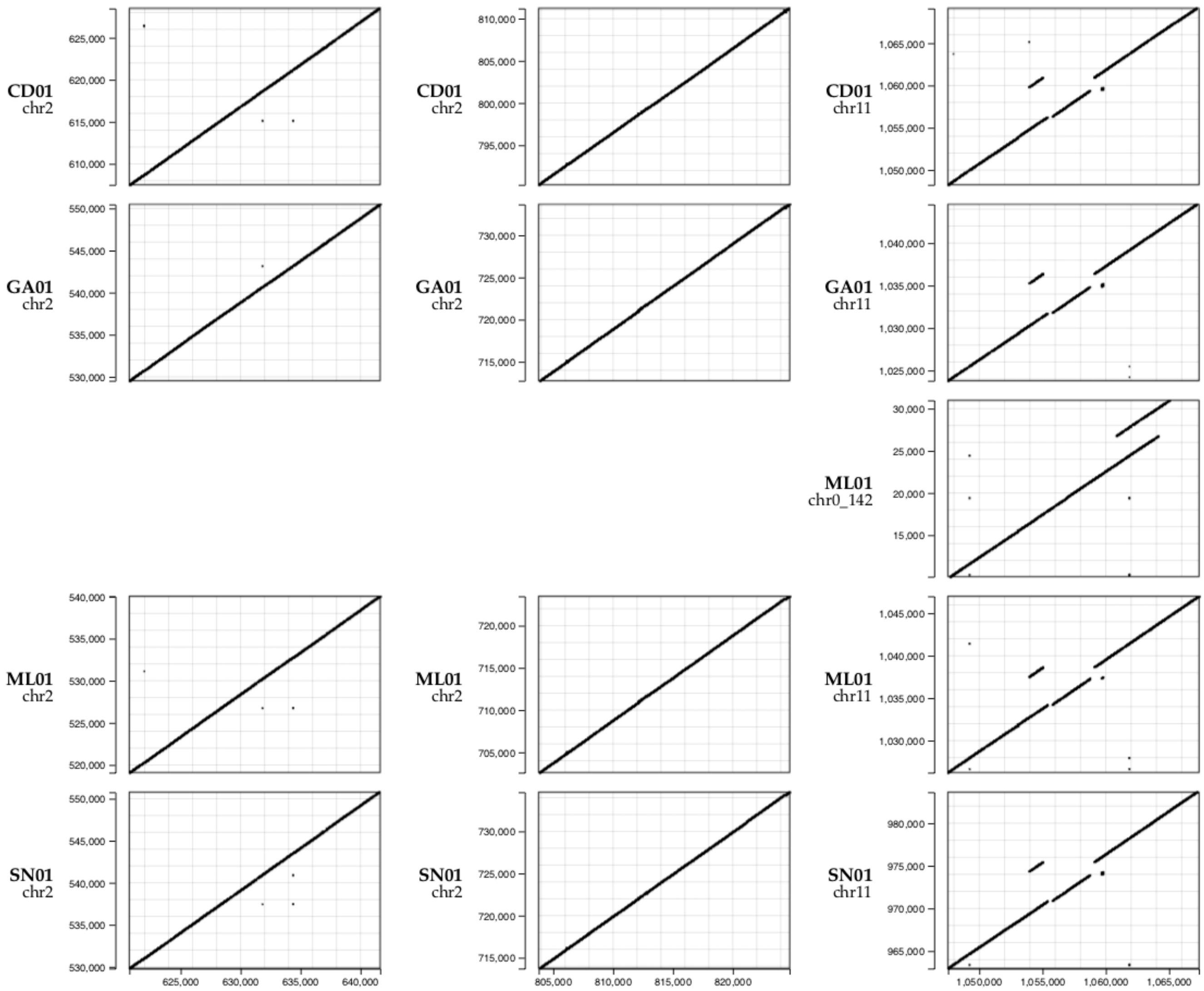

**Supplementary Figure 10. K-mer sharing between Pf3D7 and *P.falciparum* assemblies that carry *Pfsa*+ alleles.**

For each of a set of genomes of *Pf* isolates previously assembled from PacBio data [5] (rows), the plot shows short DNA sequences of length 50 (50-mers, black points) that are shared between Pf3D7 reference genome (x axis) and the specified genome assembly (y axis). Points on the diagonal indicate identical DNA structure, while off-diagonal points and breaks indicate structural variation. The *Pf* isolates selected are those that carry the HbS-associated allele at at least one of the three lead HbS-associated SNPs (as determined by aligning a 101-bp segment centred on each SNP in Pf3D7 to the corresponding assembly). In the notation of **Figure 2** the combined genotypes are: CD01 (Congo) : + + +; GA01 (Gabon): + - +; SN01 (Senegal): - - +; ML01 (Mali): - - +. Flanking sequence to the chr 11 locus aligned to two contigs in the ML01 assembly; both contigs are shown. ML01 was previously identified as containing a mixed infection[5].

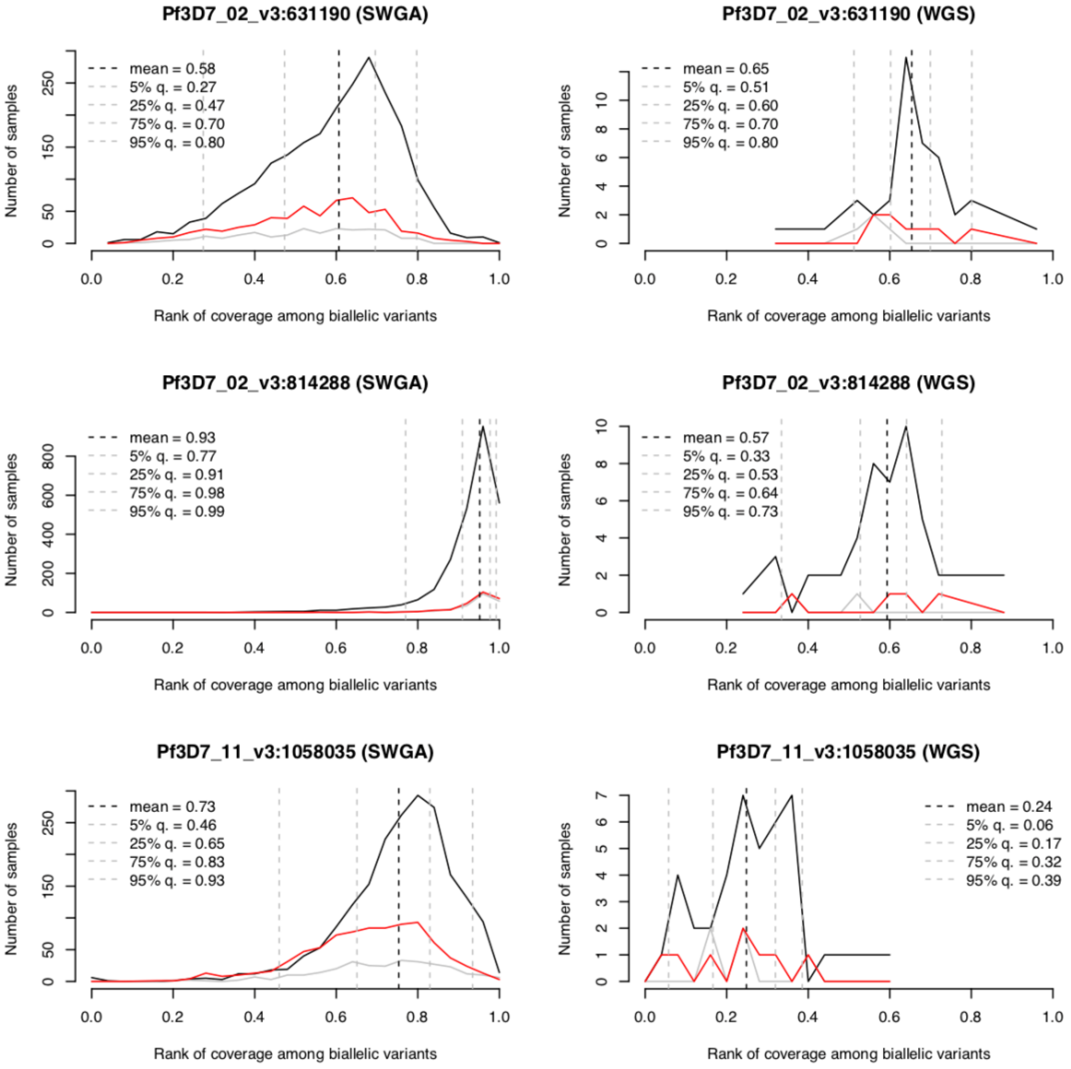

**Supplementary Figure 11. Analysis of sequence read coverage at the *Pfsa* sites.**

Panels shows normalised read coverage (x axis) against sample counts (y axis) at each of the *Pfsa1* (chr2:631,190), *Pfsa2* (chr2: 814,288) and *Pfsa3* (chr11:1,058,035) lead variants (rows), for both SWGA and whole DNA-sequenced samples. For each sample the site read coverage was normalised by computing its rank among read coverage across all biallelic sites called in our data. Results are separated by the allele carried at the focus SNP (black, reference allele; red, non-reference allele; grey, mixed genotype call). Vertical dashed lines show the mean and quantiles of the distribution of ranks.

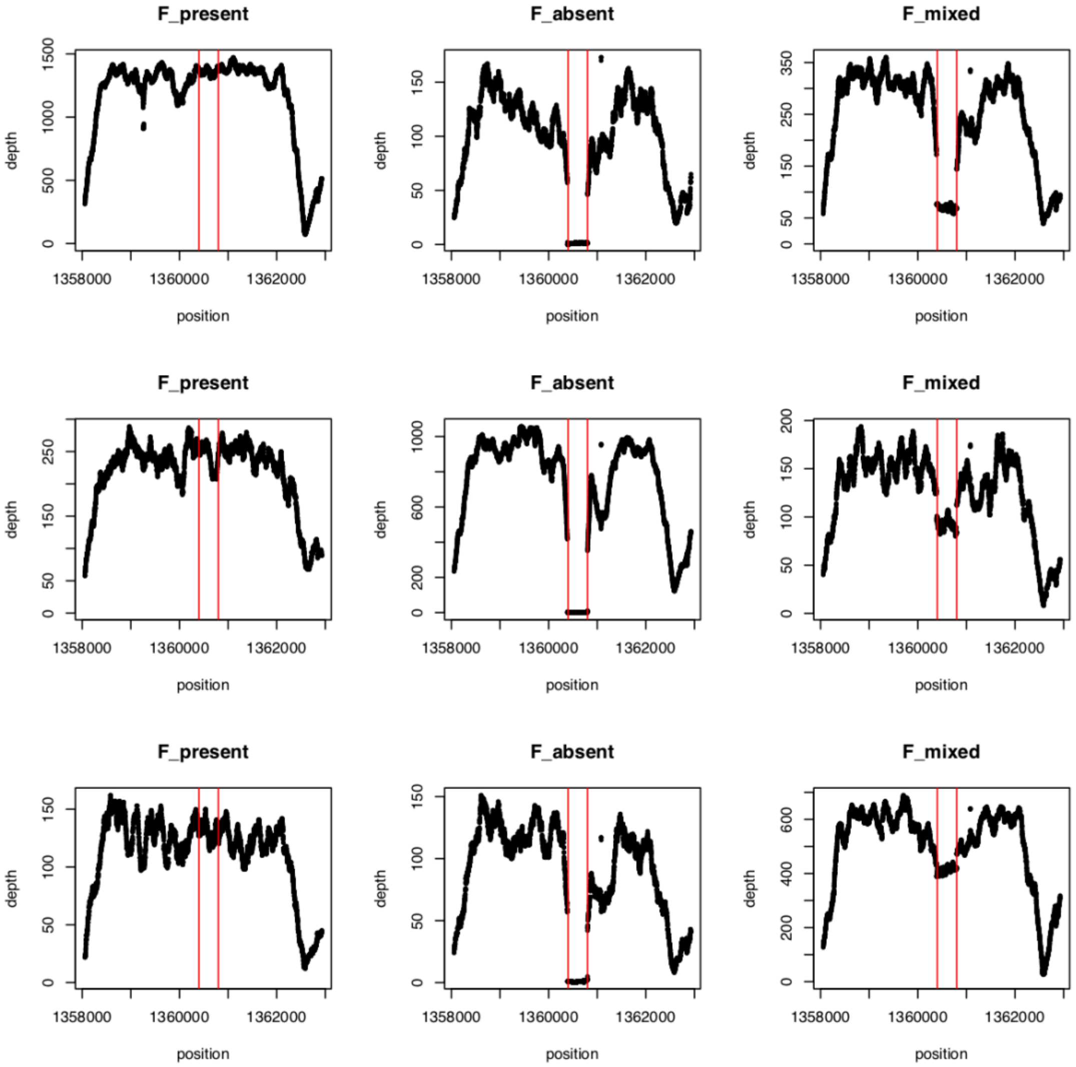

**Supplementary Figure 12. Examples of EBA175 ‘F’ segment calling.**

Panels show depth of reads aligned to the Pf3D7 genome at each site across the region of PfEBA175, for selected samples. The location of the ‘F’ segment (Pf3D7_07_v3:360,400-1,360,800) is shown between red vertical lines. The genotype called by our calling process (detailed in **Supplementary Methods**) is indicated in the panel label.
