## Supplementary Methods for "The protective effect of sickle cell haemoglobin against severe malaria depends on parasite genotype"

### 1. Supplementary Methods

#### 1.1. Building a combined dataset of human and *P.falciparum* genotypes in severe cases

##### 1.1.1. Overview

The following sections describe the sequencing and curation of a dataset of genome-wide *P.falciparum* (*Pf*) genetic variation and human genotypes from severe malaria cases collected in The Gambia and Kenya. A diagram of this process is presented in **Supplementary Figure 1**.

##### 1.1.2. Parasitaemia measurements

Parasitamia measurements were obtained for all case samples based on thin or thick blood slides at the time of sample ascertainment. Data were curated to produce a single parasitaemia measurement per individual, the count of red blood cells containing *P.falciparum* parasites per microlitre of blood (pRBCs/ul). Observed parasitaemia rates varied from very low (reported as < 10) to over $10^6$ pRBCs/ul. which represents extremely high parasitaemia (hyperparasitaemia) given typical RBC blood count of $5x10^6$ RBCs/ul.

Subsequent to selection of samples for whole DNA sequencing described below, we further linked Kenyan data to an updated set of parasitamia values re-curated from source measurements [1]. We use these values in **Supplementary Figure 2**.

#### 1.1.3. Sequencing using whole DNA samples from high-parasitaemia infections

We assessed the scope for generating coverage of the *P.falciparum* genome from sequencing whole DNA as follows. We assumed an average quantity of 40 nanograms (ng) human DNA per ul blood (based on ~5,000 white cells / ul) and that a single parasite genome weighs on the order of 25x$10^{-6}$ ng. A parasitaemia rate of $m \times 10^6$ pRBCs/ul thus corresponds to $25m$ ng/ul. This suggests sequencing whole DNA would yield a fraction of $\frac{25m}{25m+40}$ reads originating from the $Pf$ genome. For parasitaemias on the order of $10^4$ – $10^6$ pRBCs/ul we would therefore expect approximately 0.6% – 40% of reads would arise from the $Pf$ genome. For approximately 100Gbp sequencing yield and a 23Mbp genome, this suggests between 25 to over 1,000-fold $Pf$ genome coverage might be realizable using this method, depending on parasitaemia levels, although in practice we might expect this to reduce somewhat due to various forms of attrition.

Motivated by this calculation we selected a subset of case samples in each population from among those having the highest measured parasitaemia, targeting those having >$10^5$ pRBCs/ul. The Kenyan dataset sequenced also included a number of additional samples with lower parasitaemia measurements; these were selected based on their genotype at the human chromosome 4 glycophorin locus. All samples were sequenced on the Illumina XTEN platform at the Wellcome Sanger Institute. In total we obtained data for N=1,071 cases including 483 Gambians and 588 Kenyans. We aligned all reads to a combined human/parasite genome, obtained by concatenating the GRCh37 human genome reference assembly and version 3 of the Pf3D7 reference sequence [2] using BWA mem. Duplicate reads were marked with Picard MarkDuplicates and we extracted reads aligning to the $Pf$ genome for downstream analysis.

To assess $Pf$ sequencing performance we plotted coverage of the $Pf$ genome from sequencing of whole DNA against measured parasitaemia (Supplementary Figure 2). Although coverage was strongly correlated with measured parasitaemia, this correspondence was incomplete, and we noted 2 Gambian case samples with estimated zero coverage, and a larger set of 66 Gambian and 118 Kenyan cases with < 10-fold coverage. Many of these samples had relatively high measured parasitaemia and substantial coverage of the human genome (e.g. 133 with measured pRBC/ul > 10,000 and estimated > 20-fold coverage of the human genome as assessed across a region of chromosome 4). We interpret this discrepancy as resulting from a combination of noise in parasitaemia measurements as well as in possible limits to the accuracy of the curation of parasitaemia measurement data.

#### 1.1.4. Sequencing using Selective Whole Genome Amplification (SWGA)

To capture plasmodium genomes from lower-parasitaemia samples, we used Selective Whole Genome Amplification [3] (SWGA) to amplify *Pf* DNA from all cases included in our study (N=5,128 cases). SWGA was performed as previously described[3] except that we added a higher quantity (40ng) of gDNA into the reaction to allow for the mixture of parasite and human DNA. The resulting libraries were sequenced across multiple lanes on the Illumina XTEN platform at the Wellcome Sanger Institute. Reads aligning to the human reference were removed, and remaining reads from multiple lanes were merged to create sample-level read files. Remaining reads were aligned to version 3 of the Pf3D7 reference sequence using BWA mem. Duplicate reads were marked with Picard MarkDuplicates.

We plotted coverage of the *Pf* genome from SWGA sequencing against measured parasitaemia (Supplementary Figure 2). We observed relatively high coverage of the Pf3D7 genome across all reported parasitaemia levels. Sequence coverage was however variable with e.g. approximately 5% of samples having especially low coverage (258 of 5190 samples with < 10-fold coverage).

#### 1.1.5. *P.falciparum* genotype calling

To produce a robust set of parasite genotype calls we used an established pipeline that has previously been used to survey *P.falciparum* populations [4]. This pipeline uses GATK 4.0 HaplotypeCaller to identify genetic variants and to call genotypes jointly across all sequenced samples. Briefly, we first ran GATK HaplotypeCaller for each sample to generate a per-sample gVCF file, which represents a compressed view of the sequencing reads relevant for variant calling across all sites in the reference genome. We then used CombineGVCFs and GenotypeGVCFs to generate genotype calls across all samples. We specified a maximum of six alternate alleles in this process. SNPs and INDELs were then annotated using GATK's Variant Quality Score Recalibration (VQSR) based on a set of validated variants from crosses between *P. falciparum* laboratory strains [5] and based on genomic location.

Plasmodium parasites in humans have haploid genomes, but infections may consist of a mixture of parasite types due to co- or superinfection. The GATK pipeline described above calls variants as if genotypes were diploid. Heterozygous calls thus indicate regions for which reads containing both reference and non-reference alleles in substantial numbers are present (we refer to these as 'mixed' genotype calls), and homozygous calls represent variants for which the sample is largely unmixed. Although not perfect, this approach has been used previously as a practical way to handle genotype calls in mixed infections, and we adopted it here. Specifically we treated mixed genotype calls as missing data in all analyses (except where noted), and treat the remaining homozygous calls as reflecting the true haploid genotype.

#### 1.1.6. P.falciparum variants filtering

To pick a robust set of variants for analysis, we focussed on variants marked 'PASS' (i.e. with VQSLOD score > 0 and lying in the core genome which has been shown to be accessible to sequencing[5]). Inspection of remaining variants suggested the presence of many apparent multiallelic variants involving AT-rich sequence in the callset. Many of these are likely to reflect sequencing or calling errors, and we also excluded multiallelic variants from downstream analysis. In total, GATK called 4,974,562 variants across chromosomes 1-14, the apicoplast and mitochondrion, of which 2,793,802 were PASS and a subset of 1,716,459 were biallelic.

In addition to the variants above, we specifically included in our analysis variants in the region of *PfEBL1,* which is annotated as a pseudogene and lies in a subtelomeric region, but putatively encodes an erythrocyte invasion ligand in some *P.falciparum* species [6]. In total there were 1,339 biallelic variants with VQSLOD > 0 within the region Pf3D7_13_v3: 2,809,706-2,822,270.

#### 1.1.7. Generating PfEBA175 'F' segment calls

In addition to the GATK-called variants described above, we also called a known deletion variant in *PfEBA175*, which encodes an invasion ligand that binds human Glycophorin A during merozoite invasion of erythrocytes[6]. *PfEBA175* is found in two forms, the 'F' type and 'C' types, which are distinguished by the presence of one of two non-overlapping ~400 bp DNA segments. The Pf3D7 reference genome carries the F segment located between positions 1,360,400 and 1,360,800 on chromosome 7. To identify the F segment in short-read sequence data, we computed sequence coverage across each base in these segments. We considered the F segment to be present if at least 350 of the 400 sites had a coverage >=5. We considered F absent if there was evidence for unusually low coverage across this region, defined as more than 350 of the 400 sites having coverage more than 2 standard deviations below the mean, as computed at nearby single-copy sequence. If both conditions were true, a mixed genotype call was assigned, and if neither were true a missing genotype was assigned. We note that this method directly assesses the presence of the F segment, but not of the C segment which is not present in the Pf3D7 reference; however, these segments are thought to rarely if ever cooccur[7].

#### 1.1.8. P.falciparum sample filtering

To pick a robust set of samples for analysis, we filtered based on several criteria as follows. First, for samples multiply sequenced using the whole DNA and SWGA methods, we looked for discordance in genotype calls that would indicate sample mislabelling. No pair had > 2.5% discordance and we interpreted this as indicating that reads from corresponding SWGA and WGS read files represent the same DNA samples. Next, for whole DNA-sequenced samples with human genotyping available on the Illumina Omni 2.5M platform [8], we used VerifyBamId to confirm the identity of samples based on the human-aligned reads. We identified 25 samples with CHIPMIX > 2%, indicating sample contamination or sample mislabelling, and we excluded these samples (and corresponding SWGA samples) for downstream analysis. Third, we filtered samples based on the GATK v3.8.0 CallableLoci metric (defined as the proportion of reference bases where a sample has at least 5x coverage and such that at least 90% of covering reads have mapping quality >= 10, a criterion which is correlated with sequencing depth but is more directly relevant to variant calling). We excluded 758 samples where <

50% of reference bases were identified as callable by this metric, and a further 340 samples that had > 5% genotype missingness from downstream analysis. This process resulted in 4,440 samples from case individuals that passed filters.

#### **1.1.9. Curation of joint human-*Pf* analysis datasets**

For our main analysis we further restricted attention to two smaller sets of data based on the availability of human genotype data as follows. First, we formed a dataset consisting of the subset of 3,346 samples for which imputed human genotypes were previously analysed[8] ("the imputed dataset"). This contains 2,045 cases from The Gambia and 1,301 from Kenya, and contains no close relationships between human samples. We also formed a second dataset containing samples for which Sequenom MassARRAY genotyping was previously analysed[8] ("the sequenom dataset"). This set contains 4,083 samples (2,189 from The Gambia and 1,889 from Kenya) identified as not closely related based on the available genotypes[8] (although we note this determination is less certain than when using genome-wide genotype data). In total 3,246 samples were in both sets, leaving 825 with direct typing but no genome-wide data available.

### 1.2. Modelling association of host and infection genotypes

#### 1.2.1. Basic association model

In **Methods** we interpret the odds ratio computed in severe malaria cases in terms of a simplified model of infection. Here we further describe this and extend to allow for the case where parasite genotypes might evolve through the course of an infection.

As in **Methods** we let A denote a population of susceptible individuals. We assume that we are interested in a specified set of parasite variants such that parasites have one of $J+1$ possible combined genotypes, denoted $j = 0, \cdots, J$. For a given individual we write $I = x$ or more briefly $I_x$ to denote that the individual was bitten and infected with genotype x; more generally $x$ might denote a mixture of parasite genotypes. For clarity, we define "infected" here to mean that parasites are injected into the bloodstream during a bite by an infected mosquito; infections might or might not successfully further invade host cells and continue to grow. Separately, we write $G = y$ to denote that the infection genotype at the time of measurement (i.e. sample ascertainment in our study) was $y$.

In principle, the infection-time ($I$) and ascertainment-time ($G$) genotypes might differ. This could happen if the initial infection is mixed and genetic drift or selection acts on parasites within-host, or if parasites mutate during the course of infection.

We write $D$ to denote "individual has severe disease", and write $D_y$ for "individual has severe disease with parasite genotype y" (i.e. $D_y = (D, G = y)$). Lastly, $E{=}e$ denotes a particular level of a host genotype. The following table summarises this notation.

Model notation

| | |
|---|---|
| $A$ | Study population |
| $D$ | individual has severe malaria |
| $D_y$ | Individual has severe malaria with parasite genotype y |
| $I = x$ or $I_x$ | Individual was infected with parasite genotype x |
| $G = x$ | Denotes parasite genotype at time of sampling |
| $E = e$ | Individual has host genotype e |

All the probabilities we discuss are conditional on the assumed population $A$, which we drop from the notation where convenient. The odds ratio for a particular genotype $G = y$ and host genotype $E = e$ computed in severe malaria cases relative to baseline genotypes can now be written as

$$OR_{G=y,E=e} = \frac{P(D_y, E = e|D)}{P(D_0, E = e|D)} \bigg/ \frac{P(D_y, E = 0|D)}{P(D_0, E = 0|D)} \qquad \text{(S1)}$$

The observed parasite genotype depends on the infection genotype, which is unobserved, so we must sum over it:

$$\begin{aligned} P\left(D_y, E = e \middle| D\right) &= \sum_x P(D_y, I_x, E = e|D) \\ &= \frac{1}{P(D|A)} \cdot \sum_x P\left(D_y \middle| I_x, E = e\right) P(I_x, E = e) \end{aligned} \qquad \text{(S2)}$$

The first term in (S2) is independent of the genotypes and cancels out when forming the ratio (S1). Thus (S1) expands to

$$OR_{G=y,E=e} = \frac{\left(\frac{\sum_x P\left(D_y \middle| I_x, E = e\right) P(I_x, E = e)}{\sum_x P(D_0|I_x, E = e) P(I_x, E = e)}\right)}{\left(\frac{\sum_x P\left(D_y \middle| I_x, E = 0\right) P(I_x, E = 0)}{\sum_x P(D_0|I_x, E = 0) P(I_x, E = 0)}\right)} \qquad \text{(S3)}$$

#### 1.2.2. Interpretation when there is no within-host evolution

In **Methods** we make the simplifying assumption that $y = x$, that is, that genotypes do not change during the course of an infection. This is likely to be a broadly appropriate assumption for infections that are not initially mixed at the set of parasite variants of interest, since within-host mutation of specific bases is likely to be relatively rare [9] (at least until parasitaemia levels become high). In this case (S3) simplifies to

$$OR_{G=y,E=e} = \left(\frac{P\left(D \middle| I_y, E = e\right)}{P(D|I_0, E = e)} \middle/ \frac{P\left(D \middle| I_y, E = 0\right)}{P(D|I_0, E = 0)}\right) \times OR^{\text{biting}} \qquad \text{(S4)}$$

where

$$OR^{\text{biting}} = \frac{P(I_y, E = e)}{P(I_0, E = e)} \middle/ \frac{P(I_y, E = 0)}{P(I_0, E = 0)} \qquad \text{(S5)}$$

Equation (S4) is equivalent to expression (4) described in **Methods**. Since y is defined as the genotype at time of initial infection, $OR^{\text{biting}} = 1$ (for all genotypes e and y) is equivalent to statistical independence of host and parasite genotype at the time of infection. Further, if $OR^{\text{biting}} \equiv 1$, it can be shown that the first term in (S4) is one (for all genotypes *e* and *y*) if and only if host and parasite genotypes contribute multiplicatively to disease risk,

$$P\left(D \middle| I_y, E = e\right) = \mu \times \frac{P(D|I_y)}{P(D|I_0)} \times \frac{P(D|E = e)}{P(D|E = 0)} \qquad \text{(S6)}$$

where $\mu = P(D|I_0, E = 0)$ is the risk given baseline genotypes. Hence, $OR_{G=y,E=e} \neq 1$ implies either nonindependence of host and parasite genotypes at infection time or that host and parasite genotypes do not contribute multiplicatively to disease risk.

#### 1.2.3. Interpretation of the general model

The general form of (S3) allows for within-host evolution and is more complex, but we show that the analogous behaviour holds: namely, $OR_{G=y,E=e} \neq 1$ implies either nonindependence between host and infection-time genotypes, or that host and parasite genotypes do not contribute multiplicatively to disease risk, in the sense that

$$P(D_y|I_x, E = e) = \mu \times \frac{P(D_y|I_x)}{P(D_y|I_0)} \times \frac{P(D_y|E = e)}{P(D_y|E = 0)} \qquad \text{(S7)}$$

where the expressions are now extended to express the risk of disease with a specific parasite genotype y given host and (possibly different) infection genotypes. To see this, note that neither $\mu$ , $P(D_y|I_0)$, nor the last ratio in (S7) depend on the infection genotype x, and they therefore cancel out of expression (S3). If host and parasite genotypes are independent at the time of biting then also $P(I_x, E = e) = P(I_x)P(E = e)$, leading to additional cancellation, so that

$$OR_{G=y,E=e} = \frac{\frac{\sum_x P(D_y|I_x)P(I_x)}{\sum_x P(D_0|I_x)P(I_x)}}{\frac{\sum_x P(D_y|I_x)P(I_x)}{\sum_x P(D_0|I_x)P(I_x)}} = 1$$

Thus, if we rule out nonindependence at time of biting, $OR_{G=y,E=e} \neq 1$ implies deviation from the multiplicative model (S7).

#### 1.2.4. Possible causes of association

There are several mechanisms by which statistical nonindependence between host and observed parasite genotypes could arise in principle, and these make different predictions that can potentially be tested. To illustrate the distinction between these mechanisms, we consider a simplified model of infection that separates out an initial phase of within-host evolution (that produces an observable level of parasitaemia and the observable genotype *G*) and a subsequent phase of infection in which disease status is determined (but parasitaemia and genotypes do not change). As above, an infection may be mixed and if so we interpret *G* to indicate the overall proportions of genotypes making up the infection. This model can be summarized in the following diagram which shows possible causal links between variables [10]:

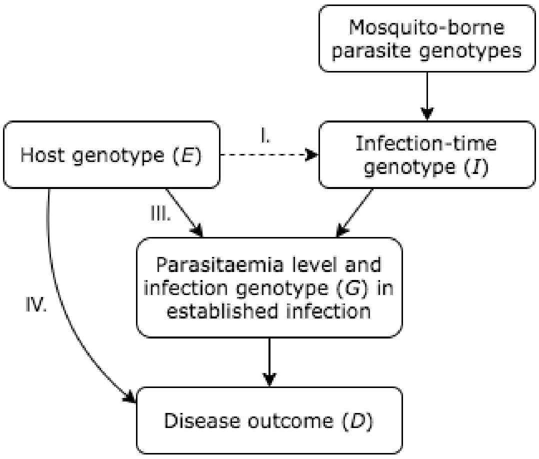

For the purposes of this illustration we ignore possible confounding factors not shown in the diagram. In this diagram there are several ways in which the odds ratio (S1) might deviate from unity as we detail below.

I. *Association induced by biting effects.* The dashed line in the diagram shows a possible influence of host genotype on parasite genotype at the time of infection. This would only be possible if genotypes of possible hosts and mosquito-borne sporozoite genotypes can detected by infectious mosquitos (or the parasites they carry).

II. *Association induced by phenotyping.* In the diagram above, the disease status *D* is a collider [10] (jointly determined by both host and parasite genotypes), and consequently conditioning on *D* could generate correlation between *E* and *I* even in the absence of a specific molecular or biological interaction. A well-known form of this is known as Berkson's paradox[11]; translating this to our setting this would occur if the clinically determined criteria for severe malaria typically arise either in infections of individuals who do not carry protective genotypes, or in infections with pathogenicity-causing alleles, or both. Specifically this leads to association when the contributions of host and parasites alleles to disease risk do not follow the multiplicative model (S6). To assess this in relation to the HbS association described in main text, we investigated whether each of the HbS-associated alleles was also associated with other known host protective alleles. Specifically we considered the protective homozygous AA genotype at rs4951377 [12,13]; genotypes carrying the G allele at rs186873296 (which tags the Dantu blood group variant DUP4) under an additive model [14,15]; and O blood group encoded by rs8176719[13]. We observed little evidence for association ($P > 0.05$ for all nine tests; P-values were not significantly divergent from a uniform distribution using a Kolmogorov-Smirnov test) although we noted that all but one estimated effect size was positive. The strongest estimated association was for rs186873296 and *Pf* chr11:1,058,035 (*OR* = 1.38; 95% CI 0.99-1.94); this estimate reduced somewhat after additionally including HbS as a predictor. Second, we also noted that the HbS-associated alleles are observed at higher frequency in community samples [4] than in the severe cases studied here (with one exception for the chr2:814,288 T allele in Gambia, which is not present in the community-sampled data but is at ~1.5% frequency in our sample of severe cases; **Figure 3**). These data do not support a formal statistical comparison due to differences in sampling, but do not appear to indicate a strong risk effect of the three HbS-associated alleles on overall disease risk. Thus, although out data do not formally rule this out, neither of these comparisons appears to support a nonspecific effect in which host protective alleles and parasite pathogenicity alleles become correlated purely due to the definition of severe disease.

III. *Host genotype effects on within-host parasite genotypes*. The most plausible explanation for association between host and parasite genotypes may be that host genotypes affect the within-host fitness of parasites, in a way that varies with parasite genotype. In the simplified model above this would lead to systematic association of host and parasite genotypes in all infections (regardless of symptom severity), but in real settings this would likely depend on the strength of selection and course of disease. For the HbS effect described in main text, this would occur if parasites with the *Pfsa+* alleles are better adapted than other parasites to growing and infecting erythrocytes of individuals with HbS genotype, compatible with the counts observed in **Figure 2**. Within-host selection of this type would likely also lead to effects on transmission of parasite genotype, and thus put selection pressure on parasite populations.

IV. *Interactions determining disease tolerance.* A separate possibility is that host and parasite genotypes jointly determine host tolerance to infection (without otherwise affecting parasite development), such that sampling of severe cases would introduce association between them. In the simplified model above, an effect of this type would be unobservable in asymptomatic cases since the effect is specific to severe disease phenotype.

### 1.3. Estimation of population relative risks using multinomial logistic regression

#### 1.3.1. Estimation using a case-population sample

We consider estimating the relative risk for severe disease observed with a particular parasite genotype *y* in population *A*,

$$RR_{E=e}(y) = \frac{P\left(D_y \middle| E = e, A\right)}{P\left(D_y \middle| E = 0, A\right)} \qquad \text{(S8)}$$

Here, as above $E=e$ denotes a particular host genotype (e.g. HbS genotype in **Figure 2**) and the relative risk is measured with respect to a chosen baseline genotype $E=0$ (i.e. non-HbS genotypes in **Figure 2**). Application of Bayes' theorem to (S8) shows that

$$RR_{E=e}(y) = \frac{P\left(E = e \middle| D_y\right)}{P\left(E = 0 \middle| D_y\right)} \Big/ \frac{P(E = e|A)}{P(E = 0|A)} \qquad \text{(S9)}$$

Expression (S9) can be recognized as an odds ratio, specifically the odds ratio comparing the frequency of the exposure *E=e* in disease cases with genotype *y* relative to the general population. It can therefore be estimated from a sample of disease cases and population controls. More generally, we show the following:

**Lemma.** *Suppose a disease with J+1 possible types* $y = 0, \cdots, J$ *follows a linear log-risk model in the population A,*

$$\log P\left(D_y \middle| E = e, Z = z, A\right) = \beta e + z^t \gamma \qquad \text{(S10)}$$

*where Z denotes a vector of covariates, and* $\beta$ *and* $\gamma$ *are log-relative risks for the exposure e and the covariates respectively. Assume for simplicity that Z consists of a single categorical covariate (i.e. z is a vector of zeros and ones with exactly one entry equal to 1). Suppose S is a case-population sample in which sampling is independent of host and parasite genotype, given the disease status and covariates. Then in the sample the multinomial logistic regression model holds:*

$$\log \frac{P\left(D_y \middle| E = e, Z = z, S\right)}{P(D_- | E = e, Z = z, S)} = \beta e + z^t \gamma' \qquad \text{(S10)}$$

*where* $D_-$ *indicates that an individual was sampled as a population control, and the coefficient* $\beta$ *of the host genotype e is the same as in the full population model.*

**Note.** This lemma is a counterpart of the well-known result that if a logistic regression model holds for a disease in the general population, then a transformed logistic regression model holds in a sample of disease cases and strict (non-diseased) controls [16]. We apply Lemma 1 in **Figure 2** to estimate the relative risk conferred by HbS on disease across multiple parasite genotypes.

**Proof of lemma.** Let Ω be the odds-ratio for disease of genotype *y* relative to the population,

$$\Omega = \frac{P\left(D_y \middle| E = e, Z = z, S\right)}{P(D_- | E = e, Z = z, S)}$$

Applying Bayes theorem to numerator and denominator gives

$$\Omega = \frac{P(E = e | D_y, Z = z, S)}{P(E = e | D_-, Z = z, S)} \cdot K_y(z) \qquad \text{(S11)}$$

where $K_y(z) = \frac{P(D_y|Z=z,S)}{P(D_0|Z=z,S)}$ is the ratio of cases of type $y$ to controls in the study. Conditional independence of sampling on the genotypes further implies that we may replace S by A in the right hand side of (S11), giving

$$\Omega = \frac{P(E = e|D_y, Z = z, A)}{P(E = e|Z = z, A)} \cdot K_y(z)$$

Applying Bayes' theorem a second time to rewrite in terms of disease risk now gives

$$\Omega = P\big(D_y\big|E = e, Z = z, A\big) \cdot \frac{K_y(z)}{\kappa_y(z)}$$

where $\kappa_y(z) = P(D_y|Z = z, A)$ is the prevalence of disease type y in the population having the given covariate levels $x$.

By assumption

$$\log\Omega = \log P\big(D_y\big|E = e, Z = z, A\big) + \log\left(\frac{K_y(z)}{\kappa_y(z)}\right)$$
$$= \beta e + z^t\gamma + \log\left(\frac{K_y(z)}{\kappa_y(z)}\right)$$
$$= \beta e + z^t\gamma'$$

for the transformed parameter $\gamma'$ defined as

$$\gamma' = \gamma + \begin{pmatrix} \log\big(K_y(z_1)/\kappa_y(z)\big) \\ \vdots \\ \log\big(K_y(z_d)/\kappa_y(z_d)\big) \end{pmatrix}$$

where $z_1, \cdots, z_d$ denote the d possible levels of the 0-1 covariate vector $z$.

### 1.4. Implementing logistic regression to test for host/parasite association

In **Methods** we describe the use of logistic regression to estimate association between host genotypes (included as predictor variables) and parasite genotypes (included as outcome variables). These results are summarized in **Supplementary Figure 4**.

Logistic regression is susceptible to finite sample bias that leads to overestimation of effect sizes in a way which varies with the frequency of predictor and outcome variables. Given that comparisons between human and parasite variants at widely varying frequencies are being tested, we chose to mitigate this by fitting the model after including a regularizing prior distribution. Following previous recommendations [17] we chose a log F(2,2) prior for this. The log-F(2,2) distribution is depicted below in comparison to a Gaussian(0, $1.87^2$) distribution (for which 95% of the mass is concentrated on a similar interval centred at zero):

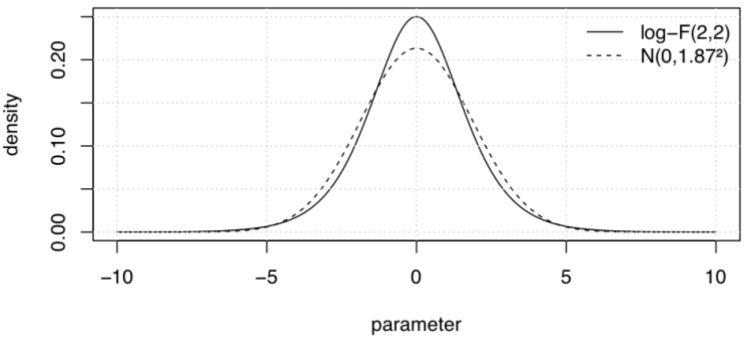

The log-F(2,2) distribution is slightly more concentrated near zero than the Gaussian (i.e. it provides greater regularization near zero) but has flatter tails (i.e. provides less regularization for parameter estimates that are far from zero). This captures an intuitive idea that most effects between individual host and parasite variants are likely to be small, but some large effects may exist.

The use of a prior enables computation of a Bayes factor for association at each pair of variants; we compute this using a Laplace approximation. A nonstandard calculation is also needed to compute a corresponding P-value; we do this by approximating the sampling distribution of the posterior mode as described in subsequent sections.

**1.4.1. Approximating the sampling distribution of the posterior mode**

We first review a standard approach that is often applied to (unregularized) regression. As sample sizes grow large the likelihood function is assumed to approach a Gaussian distribution near its maximum $v$ (up to a scaling constant which does not depend on the parameter and is ignored below),

$$P(\text{data}|\theta) \approx N(\theta|v, V) \qquad \text{(S12)}$$

Here $\theta$ denotes the vector of parameters and $V$ is a variance-covariance matrix expressing how sharply peaked the likelihood function is around is maximum. $V$ can be computed as the inverse of minus the second derivative of the log-likelihood.

As a function of the sample, the maximum likelihood estimate itself is assumed to become approximately normally distributed around the true parameter value $\theta_0$ as sample sizes grow,

$$v \sim N\left(\theta_0, \frac{I}{n}\right) \quad \text{(S13)}$$

Here *I* is another variance-covariance matrix (the Fisher information) that does not depend on the particular dataset being analysed, and *n* is the sample size. The factor of *n* in the covariance captures the fact that increasing sample size leads to estimates of increased precision, with standard errors approximately scaling as $1/\sqrt{n}$.

The standard theory links (S12) and (S13) by showing that (asymptotically as sample sizes grow large) the matrix *V* (which is estimated from the data) becomes approximately equal to $\frac{I}{n}$ (which is independent of the data) thus providing an effective way to compute P-values. Specifically, a "Wald test" P-value can be computed from the approximation $v \sim N(0, V)$ under the null model $\theta_0 = 0$; for a single component $v_i$ of $v$ this leads to

$$\text{P-value} \approx 2 \times F(-|v_i|; 0, V_i) \quad \text{(S14)}$$

where F is the cumulative distribution function of a Gaussian with the given mean and variance. (The use of $-|v_i|$ and the factor of 2 in this expression ensure that this computes a two-tailed P-value, i.e. the total mass under both tails of the distribution of parameter values greater or equal in magnitude to $v_i$).

We now consider computing P-values under regression regularized by a prior – we first consider a Gaussian prior with zero mean, i.e. defined as $\theta \sim N(0, \Sigma)$ for some variance-covariance matrix $\Sigma$. To do this, we use the same approximations as above to compute the approximate sampling distribution of the posterior mode. A Gaussian prior is particularly simple to use here because of the following well-known result which reflects the fact that the product of two gaussian densities is another gaussian density:

**Lemma.** *Under the approximation (S12), the posterior distribution is also Gaussian. Specifically,*

$$P(\theta|\text{data}) \approx N(\theta|\omega, \Omega) \quad \text{(S15)}$$

*where* $\Omega = (V^{-1} + \Sigma^{-1})^{-1}$ *and* $\omega = \Omega V^{-1} v$. *Under the approximation (S13), the posterior mode* $\omega$ *is therefore approximately distributed as*

$$\omega \sim N(\Omega V^{-1}\theta_0, \Omega V^{-1}\Omega) \quad \text{(S16)}$$

(The normalizing constant in (S15) can also be computed as another Gaussian function, which leads to the well-known computation of the approximate or asymptotic Bayes factor [18].)

It is instructive to consider these formulae in the case of a one-dimensional parameter and assuming $\theta_0 = 0$. In this case the matrices $V$ and $\Sigma$ are scalars and we have the simplifications:

$$\text{posterior mode } \omega = \frac{\Sigma}{V+\Sigma} v$$
$$\text{posterior variance-covariance } \Omega = V \cdot \left(\frac{\Sigma}{V+\Sigma}\right) \tag{S16}$$
$$\text{sampling variance of } \omega = V \cdot \left(\frac{\Sigma^2}{(V+\Sigma)^2}\right)$$

Thus the posterior mode $\omega$ is closer to zero than the maximum likelihood estimate $v$ (i.e. it is a "shrinkage estimate"); the posterior variance $\Omega$ is smaller than the likelihood variance V; and the sampling variance of $\omega$ is smaller still. The degree of shrinkage in each case depends on the relative magnitude of the likelihood variance V and the prior variance $\Sigma$, with the extremes being $\Sigma = \infty$ (which produces no shrinkage at all) and $\Sigma = 0$ (which makes all three expressions equal to zero).

In our implementation, for each parasite variant and each human variant considered, we obtain the posterior mode $\omega$ by numerical approximation using a modified Newton-Raphson with line search [19]. We then compute the approximate posterior variance-covariance matrix $\hat{\Omega}$ (computed as the inverse of negative the second derivative of the log-posterior at $\omega$) and an approximate likelihood covariance $\hat{V}$ (computed as the inverse of negative the second derivative of the log-likelihood at $\omega$). By analogy with (S14) we then compute a P-value for the parameter of interest $\omega_i$ as

$$\text{P-value} = 2 \times F\left(-|\omega_i|; 0, \left[\hat{\Omega}\hat{V}^{-1}\hat{\Omega}\right]_i\right) \tag{S17}$$

#### 1.4.2. Implementation using a log-F prior

As described above, in our implementation we chose to use the log F distribution which is a natural choice for logistic regression problems[17] and provides slightly stronger regularization near zero than a Gaussian with similar tails. There are therefore two ways in which (S17) might fail to give an accurate P-value. First, if the asymptotic approximations (S12) and (S13) fail then (S14) may not be accurate; in this case (S17) and (S14) might also differ. Second, the expressions might differ because of differences between the log-F prior and the Gaussian. To assess the impact of these, we conducted a simulation study as follows

1. We considered human and parasite variants at population frequencies of 1%, 5%, 10%, 20%, 30% and 50%.
2. For each frequency *f*, we simulated genotypes for 10 human variants in *N=3,346* samples by binomial sampling given the frequency.
3. For each frequency *f*, we also simulated genotypes for 10,000 parasite variants in *N=3,346* samples by binomial sampling given the frequency (thus representing no association between human and parasite genotypes).

4. We ran hptest to test for association between each human and parasite variant (3.6 million tests in total) with no prior applied and computed a Wald test P-value (S14) and a likelihood ratio test P-value.
5. We ran hptest a second time applying the log-F(2,2) prior and applied (S17) to recompute the P-value.
6. We plotted results stratified by the minimum count observed across all combined host and parasite genotypes at the two variants (i.e. the minimum value in the 2x2 contigency table formed by the two genotypes).

The following image shows a comparison of Wald test P-values and parameter estimates for the unregularized and log-F(2,2)-regularised regression.

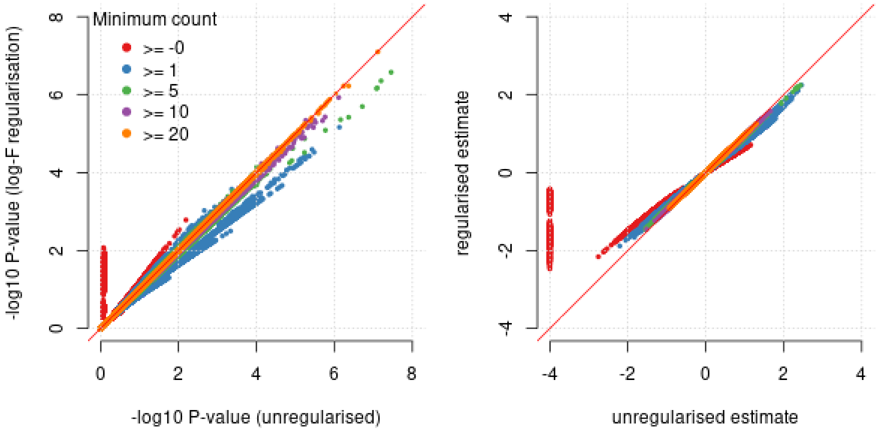

We noted that when the minimum combined genotype count is at least 20, the P-values computed by (S14) and (S17) are essentially identical, although discrepancies can be observed for lower counts; the P-value computed from regularised regression is typically more conservative in these cases. These discrepancies are similar in magnitude to those observed between Wald and likelihood ratio test P-values computed from unregularized regression. For smaller counts, inclusion of the prior has the desirable property that it generates less overestimation of effect size magnitude, including for some pairs of variants that generate extremely large estimates when the prior is not included.

### 1.5. Investigation of additional signals of association

#### 1.5.1. Overall interpretation of additional signals

In addition to the HbS associations described in main text, we observed additional candidate associations between other human and *Pf* variants in our discovery data. The evidence for these associations is likely not strong enough to establish these associations without additional information. A full list of associations can be found in **Supplementary Table 1**; we detail those with $BF > 10^5$ and those involving other = previously established human protective mutations at the ABO, ATP2B4 and glycophorin regions below. Additionally, we also observed a larger number of *Pf* variants associated at lower levels of evidence ($BF > 10^3$) with HbS and we interpret these below.

#### 1.5.2. Association between *GCNT2* and two regions of the *Pf* genome

We observed a candidate host-parasite genetic association intronic variation in the gene *GCNT2* (lead SNP: rs517371 chr6:10,554,048 C>G) and a non-synonymous SNP in MSP4 (*Pf* chr2:278,302 T>C). (*BF*=$2.8 \times 10^{6}$; *P* = $1.4 \times 10^{-9}$; *OR* = 0.39 (0.28-0.53) for effect of human 'G' allele on parasite 'C' allele; **Supplementary Table 1**). *GCNT2* determines the two reciprocal antigens of the I blood group by adding a β1,6-linked polylactosamine side chain onto the i antigen to generate a branched I antigen. Besides newborns and individuals with rare inactivating mutations in *GCNT2*, all individuals express both I and i antigens on the erythrocyte surface to varying degrees, with I expression dominating [20]. *GCNT2* has three alternative first exons that have cell-type specific expression. The associated variants lie just upstream of the 2nd first exon that is expressed in multiple cell types, but not in erythrocytes which are thought to use the 3rd first exon isoform [21]. It is thus unclear whether these variants affect gene or transcript expression in relevant cell types. P. falciparum's MSP proteins are generally thought to act in early stages of RBC invasion, but the specific function of MSP4 remains unknown [6].

Interestingly, an additional signal of association between *GCNT2* variation (rs78972384 C > T) and *Pf* variation in *RH1* (chr:138,623 A > T) was also observed (*BF*=4.9; $1.5 \times 10^{-6}$). We are unaware of any existing evidence for a molecular interaction between the I antigen with *PfMSP4* or *PfRH1*, or with parasite invasion more generally.

#### 1.5.3. Association between HLA alleles and variation in several regions of the *Pf* genome

As shown in **Supplementary Table 1**, a number of HLA alleles associate with variation in the parasite genome with Bayes factors in the range $10^{5} – 10^{6}$. These include including HLA-A*68 (associated with variation in *PfWDTC1* "WD and tetratricopeptide repeats protein 1, putative"); HLA-B*49 (associated with variation in *PF3D7_1141700* "OTU domain-containing protein, putative"); HLA-DQB1*03 (associated with variation in *PF3D7_0714900* "tRNA Serine"); HLA-A*01 (associated with variation in *PfXL2* "Exported lipase 2"); and HLA-DPA*02 (associated with variation in *PfSET3* "SET domain protein, putative"). We caution that HLA alleles in our data were obtained by imputation using a reference panels with limited coverage of African populations[8]; however, the alleles listed here have reasonably allele frequency ( > 4.5% in both countries) and reasonably imputation confidence (IMPUTE info > 0.94 in both countries).

Among these candidate associations, the association between HLA-A*01 and *Pf* chr10:86,025 A>T may be notable because the *Pf* 'A' allele was only observed in infections of individuals that do not carry the HLA-A*01 allele, leading to a large estimated effect size (OR= 0.06).

#### 1.5.4. Association with ABO, ATP2B4 and glycophorin variation

We also note here weak evidence in our discovery data (BF > $10^{3}$) for association between previously established host protective mutations other than HbS[8] and *Pf* variation. We observed weak evidence for association of the O blood group mutation (rs8176719) and an intergenic variant near *PfAROM* (Pf chr2:257614 C>G; BF = $1.6 \times 10^{4}$). The *ATP2B4* variant rs4951377 was associated with an intergenic variant on *Pf* chromosome 7 (chr7: 507,739 G > A; BF = $7.3 \times 10^3$). The glycophorin structural variant DUP4 was not associated with any variants with BF > $10^3$.

#### 1.5.5. Additional associations with HbS

In addition to the *Pfsa1*, *Pfsa2* and *Pfsa3* loci described in main text, a number of other *Pf* loci appear to be associated with HbS at lower levels of evidence (*BF* > $10^3$; **Supplementary Table 1**). These include missense variants in both *PfCLAG3.1* and *PfCLAG3.2* (cytoadherence-linked asexual protein 3) on chromosome 3, in *PfLSA1* (Liver-stage antigen 1), a noncoding variant close to *PfREX2* (Ring-exported protein 2) and variation in *PfEBL1* (erythrocyte binding like protein 1). These variants are also among those observed to be in strong LD with *Pfsa* variants (**Supplementary Table 3**) and we interpret them as reflecting the same association signal.
